# Mapping the Human Proteome with Physical Access to DNA

**DOI:** 10.1101/2024.04.04.588092

**Authors:** Jakob Trendel, Simon Trendel, Shuyao Sha, Bernhard Küster

## Abstract

In a human cell, DNA is packed in histones, RNA, and chromatin-associated proteins, forming a cohesive gel. At any given moment, only a specific subset of the proteome has physical access to the DNA and organizes its structure, transcription, replication, repair and other molecular functions essential to the way the genome is read and maintained. We have developed a ‘zero-distance’ photo-crosslinking approach to quantify proteins in direct contact with DNA in living cells. Collecting DNA interactomes from human breast cancer cells, we present an atlas of over one thousand proteins with physical access to DNA, and hundreds of peptide-nucleotide crosslinks pinpointing protein-DNA interfaces with single amino-acid resolution. Differential comparisons of DNA interactomes from cells undergoing treatment with estrogen or genotoxic chemotherapy recapitulated the recruitment of key transcription factors and DNA damage proteins. This opens a direct way to explore genomic regulation in a hypothesis-free manner, applicable to many organisms and systems.

## Introduction

What proteins interact with the genome determines the fate of a cell and often the fate of entire organisms (Lambert et al., 2018). DNA is embedded in a dense meshwork of protein and RNA generally referred to as ‘chromatin’, where protein-DNA interactions can occur directly, as for example in the case of histones, or indirectly, as in the case histone-modifying complexes (Bannister and Kouzarides, 2011). For transcription factors it has been demonstrated that direct interactions with DNA can have a different effect than indirect DNA interactions within complexes (Avsec et al., 2021; Gordân et al., 2009). The overall composition of the subproteome with physical access to DNA is momentarily ill-defined. Proteomic assessments of the DNA interactome has so far been obscured by the gel-like nature of chromatin (Strickfaden et al., 2020), whose boundaries are fuzzy and usually defined by the method of fixation and isolation (Kustatscher et al., 2014a). Previous studies interrogating chromatin composition in human cells have reported anywhere between 1500-3500 proteins (Ginno et al., 2018; Kustatscher et al., 2014b; Rafiee et al., 2021). While condensed mitotic chromosomes can be purified for proteomic analysis because they form stable complexes (Lewis and Laemmli, 1982; Ohta et al., 2010), covalent crosslinking is required to fix protein-DNA interactions of interphase chromosomes because their DNA is dispersed in the nucleoplasm and therefore cannot be purified as one physical entity. Proteins can be covalently crosslinked to nucleic acids by chemical crosslinking, typically with formaldehyde (FA), or by ultra violet (UV) light-induced activation of the natural DNA bases. FA crosslinking is highly effective for covalently fixing protein-DNA interactions, however, it also leads to indirect crosslinking of proteins with other proteins and RNA within the chromatin gel. This makes it impossible to differentiate direct from indirect interactions and creates a tripartite molecule that is notoriously hard to purify for proteomic analysis (Ginno et al., 2018; Kustatscher et al., 2014b; Rafiee et al., 2021). In contrast, photo-crosslinking only creates covalent crosslinks between UV-activated nucleic acid bases and proteins physically interacting with them, and has therefore been extensively applied for the proteomic interrogation of the RNA-bound proteome (Baltz et al., 2012; Castello et al., 2012; Gebauer et al., 2020; Hentze et al., 2018; Queiroz et al., 2019; Trendel et al., 2019). Photo-crosslinking of DNA, however, has been challenging due to the lower photo-reactivity of DNA compared to RNA (Smith, 1969; Smith and Meun, 1968). Proteomic studies that isolated UV-crosslinked DNA-peptide hybrids were so far able to report six photo-crosslinks mapping to ten highly abundant proteins from intact mouse embryonic stem cells using a 258 nm laser (Reim et al., 2020), and 36 photo-crosslinks mapping to 27 proteins from HeLa nuclei using conventional 254 nm bulbs (Stützer et al., 2020). Other studies that enriched entire proteins photo-crosslinked to DNA for their proteomic analysis tried to increase the yield of the crosslinking reaction by metabolically labeling the DNA of proliferating cells with the photo-activatable deoxyribonucleotide 6-thioguanine prior to irradiation (Gueranger et al., 2011; Guven et al., 2016). Similar to proteomic approaches using chemical crosslinking, these studies achieved only poor elimination of non-crosslinked proteins during the workup of photo-crosslinked protein-DNA complexes, resulting in high proteomic background and strongly suppressed dynamic range when quantitatively comparing samples (Ginno et al., 2018; Guven et al., 2016). Thus, while the systematic quantification of direct protein interactions with the genome is highly desirable (Van Mierlo and Vermeulen, 2021), there is currently no adequate methodology to i) effectively photo-crosslink protein in physical contact with DNA, and ii) to extract protein-crosslinked DNA with high purity.

Here, we present an optimized photo-crosslinking approach for interrogating direct protein-DNA interactions in living cells, addressing these challenges. Therefore, we combined metabolic DNA labeling of cultured cells with the photo-activatable 4-thiothymidine (4ST) and a specialized high-intensity 365 nm LED-based irradiation system. For the proteomic analysis of photo-crosslinked protein-DNA complexes by liquid-chromatography and mass spectrometry (LC-MS), we developed a multistep process of denaturing purifications named XDNAX for ‘protein-crosslinked DNA extraction’, which strongly enriched more than 1800 proteins photo-crosslinked to DNA while eliminating most non-crosslinked proteins below the limit of detection. Serendipitously, we identified numerous nucleotide-crosslinked peptides in XDNAX samples without further enrichment, which we employed as additional evidence to characterize the protein-DNA interface. Across all presented experiments we found 635 nucleotide-crosslinked peptides mapping to 358 proteins from intact MCF7 cells – a ten-fold increase over previous reports (Reim et al., 2020; Stützer et al., 2020). By quantifying DNA interactomes differentially between treatments, we show that our photo-crosslinking approach can identify transcription factors responding to cellular perturbations and discover vulnerabilities in the DNA repair machinery of breast cancer cells treated with different genotoxic chemotherapeutics. Overall, the presented DNA interactomes provide in-cell insights on direct protein-DNA interfaces, genomic regulation, and the mechanism of genotoxic drugs.

## Results

### A high-powered UV Irradiation System for the activation of photo-reactive nucleotides in living cells

To catalogue and quantify human proteins accessing the genome, we devised a photo-crosslinking procedure that efficiently couples cellular DNA of living cells to the proteins in its direct proximity. Previous studies using UV light in the range of 254 nm (UVC) for the photo-activation of natural DNA bases detected very few DNA-crosslinked proteins, indicating that higher light intensities might be required. However, high intensity UVC light can cause unwanted fragmentation of DNA and protein molecules. To increase crosslinking yields while avoiding photodamage, we decided to apply 365 nm light (UVA) in combination with metabolic labeling of the DNA with 4-thiothymidine (4ST). Unlike naturally occurring nucleic acids, 4ST is photo-activated at higher wavelengths. Figure S1A shows that this circumvented light scavenging of cellular macromolecules, presumably preventing photodamage and fragmentation of the natural DNA, which we intended to use as a purification handle in subsequent sample processing steps. Because of their application in UV curing of resins and glues, very powerful 365 nm UV-LEDs have become commercially available. Their UV emission rival light intensities of lasers despite much larger fields of irradiation. While 365 nm light is not ideal for photoactivation of 4ST, which has an absorption maximum of 330 nM (Figure S1A), we anticipated that the enormous photon output of 365 nm LEDs would compensate for the approximately 90 % poorer light absorption at this wavelength offset. Notably, LEDs with lower emission wavelengths do exist, yet, they currently exhibit much inferior intensities of less than 1 % of the most powerful 365 nM LEDs. We constructed an irradiation device that employs an array of high-intensity 365 nM LEDs, able to evenly irradiate cell culture dishes of 15 cm in diameter (Figure 1A & Figure 1B). The field of irradiation was more than 600 times larger than previous laser-based irradiation systems at very similar light intensity (∼0.3 cm^2^ for a 258 nm UV-laser system (Steube et al., 2017) vs. ∼175 cm^2^ for our 365 nm UV LED system, both at ∼2000 mW/cm^2^). In comparison to a conventional, bulb-based UV irradiation device, our system was able to emit three orders of magnitude more energy (Figure 1C). By combining two independent cooling systems for both the LEDs and the irradiation chamber, cells could be covered by PBS or cold medium and maintained at four degrees Celsius throughout the irradiation process. We named the device UVEN for ‘UV irradiation system for ENhanced photo-activation in living cells’ and benchmarked it for photo-crosslinking by irradiating aqueous solutions of 4ST or 4-thiouridine (4SU), which is commonly applied to enhance protein-RNA crosslinking in living cells (Baltz et al., 2012; Hafner et al., 2010). Figure 1D shows that the absorbance of either nucleotide decreased upon prolonged irradiation, whereas no change was seen when they were irradiated with a conventional UV bulb (see also Figures S1B & Figure S1C). Using mass spectrometry, fast formation of compounds with masses corresponding to previously described photo-dimers (Massey et al., 2001) was observed in the UVEN-irradiated samples, while the formation was much slower under bulb irradiation (Figure 1E & Figure S1D). This confirmed that UVEN effectively photo-activated thiol-containing nucleobases despite its imperfect emission wavelength. It also indicated that UVEN accelerated the photo-activation approximately 100-fold in comparison to a standard device. Analogously to naturally occurring nucleotides, we found the ribonucleotide 4SU about ten times more reactive than the deoxyribonucleotide 4ST, explaining why the use of 4ST for crosslinking proteomics has been very limited so far. Nevertheless, metabolic labeling of expanding cancer cells with 4ST and its photoactivation has been explored for photodynamic therapy in cell culture and a rat xenograft model, where UVA irradiation killed 4ST-labeled cells but not unlabeled cells (Massey et al., 2001; Pridgeon et al., 2011; Reelfs et al., 2007). We exploited this known photo-sensitization of cells by 4ST to demonstrate that UVEN could activate 4ST in living cells and to determine the optimal irradiation time. Therefore, MCF7 cells were expanded in the presence of increasing 4ST concentrations and their proliferation monitored by live cell imaging (Figure 1F). After four days cells were irradiated with UVEN and their proliferation further recorded in fresh media without 4ST. At 100 µM 4ST labeling already six seconds of irradiation were enough to completely halt cell expansion. Unlabeled control cells, however, continued proliferation even after 60 seconds of UVEN irradiation, indicating that i) natural DNA remained mostly unharmed by light at this wavelength and ii) cells were alive throughout the irradiation. We investigated what concentrations of 4ST would be tolerated by cultured cell and compared the proliferation of MCF7 (breast adenocarcinoma) and U2OS cells (epithelial osteosarcoma) in the presence of increasing doses of 4ST or 4SU. With a half inhibitory concentration (IC_50_) between 1-2.5 mM, 4ST was much better tolerated than 4SU with an IC_50_ between 100-200 µM (Figure S1E). For our subsequent photo-crosslinking experiments, we chose a concentration of 100 µM 4ST for labeling cells before one minute of UVEN irradiation in ice-cold PBS, with the intention to ‘freeze’ protein-DNA interactions during irradiation and suppress cellular responses to the emerging crosslinks.

**Figure 1:**
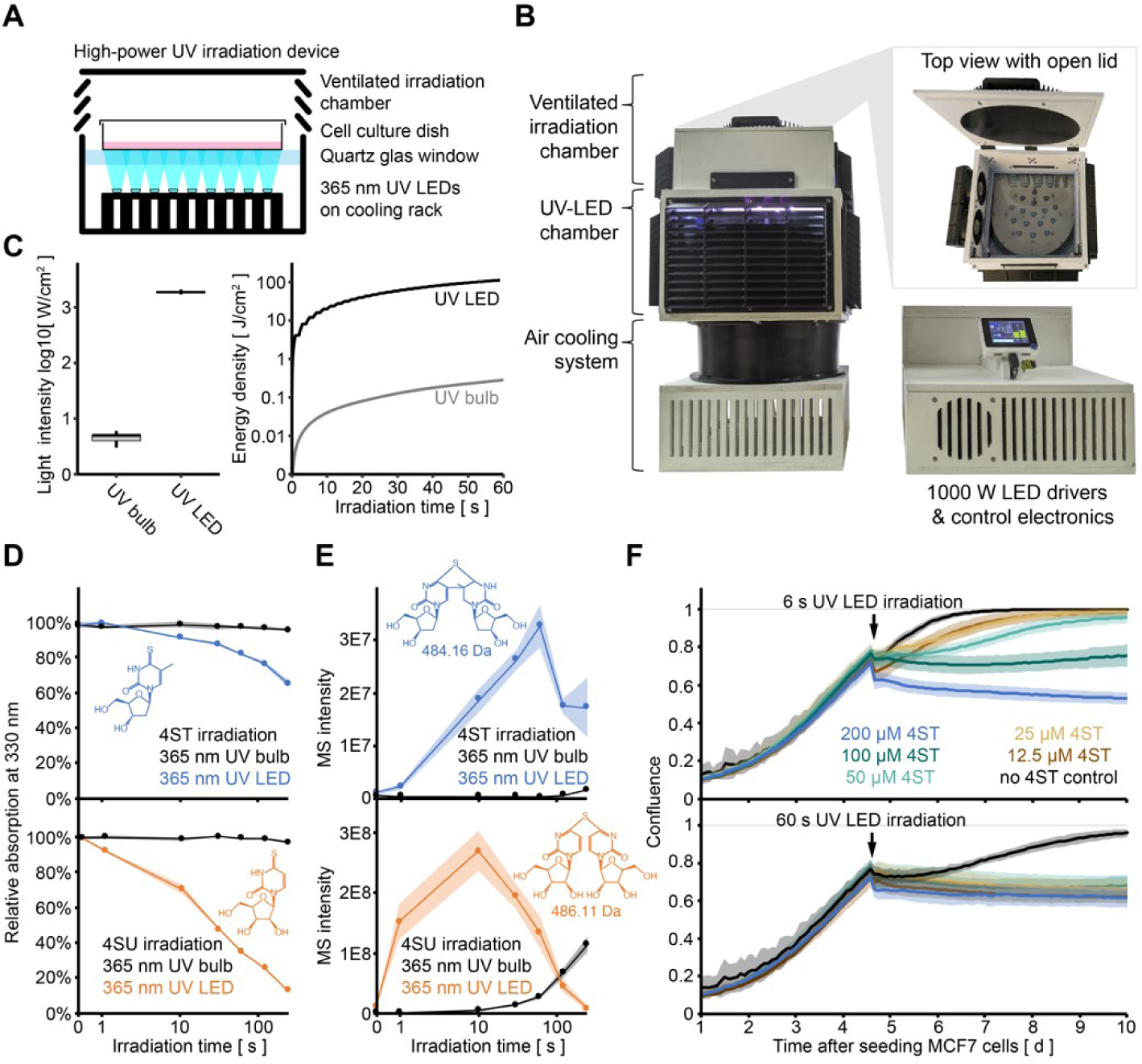
Photo-activation of the UV-reactive deoxyribonucleotide 4-thiothymidine in living cells. A) Schematic representation of the high-intensity UV-irradiation device UVEN. Cell culture dishes of up to 15 cm diameter are irradiated from below by high-powered 365 nm UV-LEDs. B) Photos of the actual UV-irradiation device UVEN. The irradiation tower (left) contains the irradiation chamber that can be filled from above (insert) on top of the LED chamber that is cooled from below. Current drivers and control electronics are located in a separate housing (right). C) Comparison of UV emissions by a conventional 365 nm UV bulb-based irradiation device (UV bulb) and UVEN (UV LED). In both cases a UV photometer was placed in 3.5 cm distance to the light source to record light emission between 340-405 nm wavelength. D) Time series displaying 330 nm absorption measurements of 4-thiothymidine (4ST, top) or 4-thiouridine (4SU, bottom) irradiated in water for the indicated time (note logarithmic scaling). High-power UVEN irradiation (blue/orange) is compared to conventional bulb irradiation (black). Displayed are means of triplicates with shaded areas indicating one standard deviation. See also Figure S1B & Figure S1C. E) Time series showing the formation of 4ST (top) or 4SU (bottom) photo-dimers upon irradiation in water (see Figure S1D) as detected by LC-MS. Displayed are integrated ion intensities for the indicated masses for the same samples as in D. F) Lineplot showing the confluence of MCF7 cells monitored by live cell imaging. Cells were grown in the presence of the indicated concentration of 4ST, irradiated for 6 s (top) or 60 s (bottom), and returned to culture in fresh medium.

### Extraction of protein-crosslinked DNA from photo-sensitized cultured cells

In order to extract photo-crosslinked protein-DNA complexes for their analysis by LC-MS we devised a denaturing purification procedure aimed at eliminating all non-crosslinked protein from DNA, yielding only DNA-crosslinked proteins with direct access to the genome at the moment of irradiation (Figure 2A). During conventional thiocyanate-phenol-chloroform (TRIZOL) extraction, chromatin will collect in the interphase and can be recovered as cohesive film, removing much of the non-crosslinked proteins, RNA, lipids and other cellular debris concentrated in the organic and aqueous phases (Chomczynski and Sacchi, 2006; Trendel et al., 2019). We treated the interphase with an additional iteration of the TRIZOL procedure to remove the excess of trapped, non-crosslinked protein (Queiroz et al., 2019), and then resolubilized it for RNase digestion. The resulting extract of protein-crosslinked DNA was sheared by ultrasonication, denatured by boiling in guanidinium thiocyanate and vigorously washed on silica spin columns to remove non-crosslinked protein to completion. Photo-crosslinked protein was released from DNA by nuclease treatment and trypsin digested for LC-MS analysis. We named this method ‘extraction of protein-crosslinked DNA’ or XDNAX (for details see Methods).

**Figure 2:**
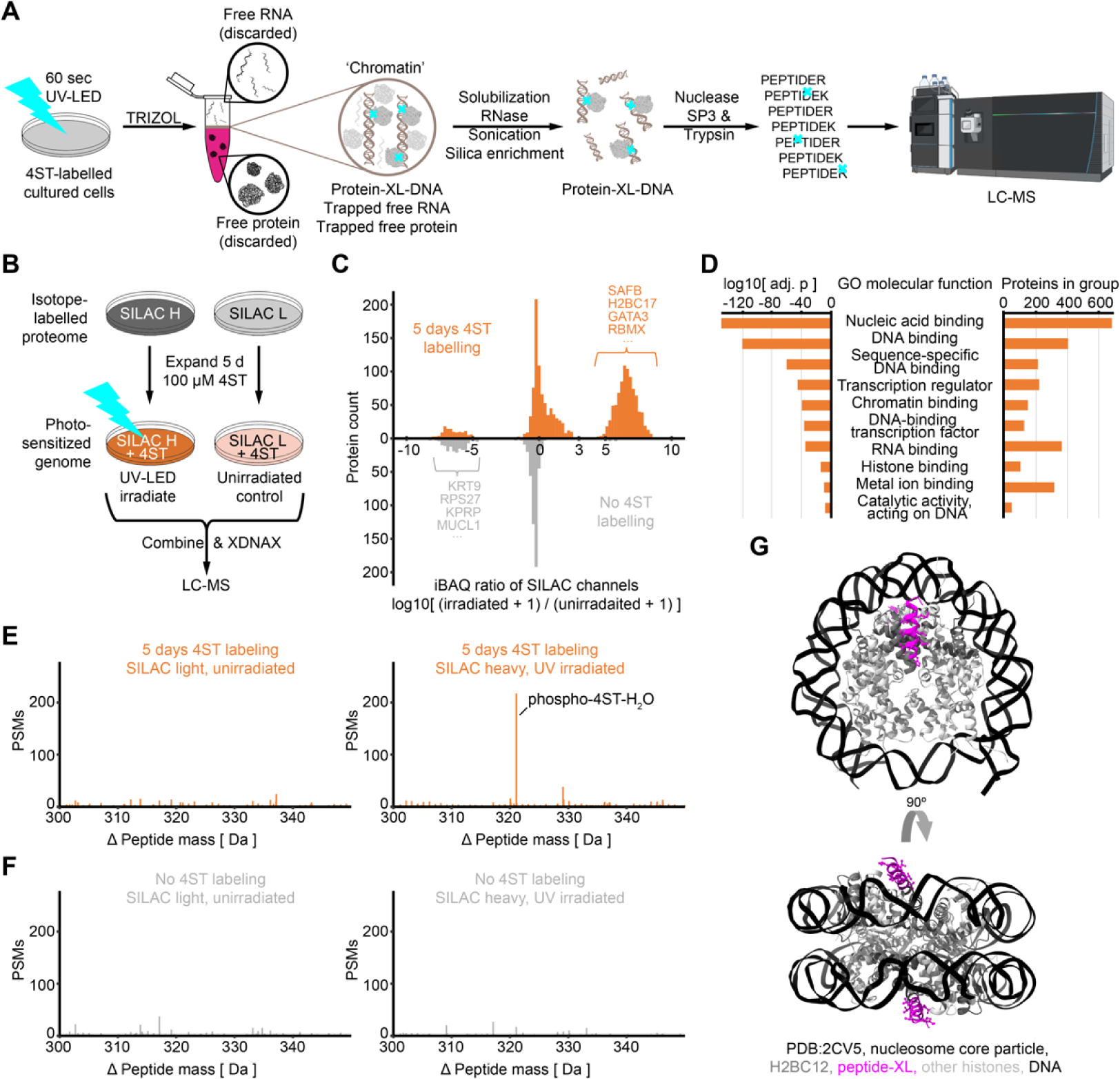
Extraction and proteomic analysis of protein photo-crosslinking to DNA of live cells. A) Workflow for the extraction of protein-crosslinked DNA or XDNAX. After photo-crosslinking cells are lysed in TRIZOL and chromatin recovered from the interphase. The DNA is then purified from non-crosslinked protein by silica column enrichment and crosslinked DNA-protein complexes treated with nuclease and trypsin before quantification by LC-MS.| B) Experimental scheme for the validation of XDNAX using SILAC. For details see text. C) Histogram illustrating the enrichment of photo-crosslinked protein extracted by XDNAX from MCF7 cells according to B (orange, top). Because most proteins did not show any intensity in the unirradiated SILAC channel iBAQ ratios are used and pseudo counts added to each SILAC channel. As control the outcome of the experiment using cells without prior 4ST photo-sensitization is shown mirrored below (grey, bottom). D) Bar plot showing the enrichment and protein group size of ten selected GO terms. All proteins in the heavy SILAC channel from C were compared to a deep MCF7 full proteome. E) Histogram of peptide-spectrum matches (PSMs) identified in an open search on the data in C by the mass-tolerant search engine MSfragger. For visibility only a Δ mass window around the 4ST modification is shown, for the full Δ mass range see Figures S2B and S2C. F) Same as in E but for control experiment omitting 4ST labeling. G) Crystal structure of the human core particle (PDB 2CV5) illustrating the position of the nucleotide-crosslinked peptide from histone H2BC12 identified in our DNA interactome (magenta) relative to the DNA (black).

The enrichment of photo-crosslinked protein via XDNAX was evaluated by the experiment displayed in Figure 2B, where stable isotope labeling in cell culture (SILAC) (Ong et al., 2002) was applied to compare two populations of MCF7 cells, which were both initially expanded for five days in the presence of 4ST. Subsequently, SILAC heavy cells were irradiated with UVEN, whereas SILAC light cells were left unirradiated. Cells were combined and subjected to XDNAX so that the MS signal in the heavy SILAC channel would indicate protein enriched upon photo-crosslinking while signal in the light SILAC channel would indicate non-crosslinked background. Indeed, our optimized workflow led to very strong enrichment of protein from the irradiated cells so that 1026 of the 1803 quantified proteins were enriched by more than ten-fold and 876 proteins were exclusively detected in the SILAC channel corresponding to irradiated cells (Figure 2C, top). Moreover, when we performed the same experiment on cells whose genome was not photo-sensitized with 4ST, we did not observe any enrichment (Figure 2C, bottom). Gene ontology (GO) analysis comparing proteins exclusively found in the irradiated SILAC channel after XDNAX against a deep full proteome of the same cell line (>10000 proteins, see Methods for details) confirmed strong enrichment for proteins annotated as nucleic-acid binding (p=1E-149) and more specifically DNA-binding (p=1E-120) (Figure 2D). As expected, intensity-based quantification of this SILAC channel showed especially histones among the most abundant proteins, followed by splicing factors and chromatin-organizing proteins (Figure S2A). In order to investigate the preservation of post-translational modifications on the enriched proteins we applied the mass-tolerant search engine MSfragger and found phosphorylation as well as ubiquitination well conserved after XDNAX (Figure S2B). To our great interest, we found an additional modification of 321 Da occurring only in the irradiated SILAC channel (Figure 2E, Figure S2B). The modification did not occur when 4ST-labeling was omitted (Figure 2F, Figure S2C) and corresponded exactly to the mass of phosphorylated 4ST minus a water molecule, a loss previously observed for uridine-crosslinked peptides in the context of protein-RNA crosslinking (Kramer et al., 2014; Trendel et al., 2019). We highlight this observation because it directly confirms covalent crosslinking between protein and DNA and shows the powerful enrichment achieved by XDNAX. Notably, no additional purification step was necessary for the identification of these nucleotide-crosslinked peptides, which is markedly different from previous studies investigating peptide-RNA or peptide-DNA crosslinks that required elimination of non-crosslinked peptides (Bae et al., 2020; Kramer et al., 2014; Reim et al., 2020; Stützer et al., 2020; Trendel et al., 2019). Two of the most frequently observed hybrids were derived from the core histone H2BC12 (Figure 2G) and the structural protein HMGB1 (Figure S2D), for which co-crystal structures complexed with DNA were available that illustrated the interface between nucleotide-crosslinked peptide and DNA.

### An atlas of proteins with direct access to DNA

In order to take inventory of proteins directly interacting with DNA, we applied our photo-crosslinking approach to five replicates of MCF7 cells from which we derived a catalogue of direct DNA interactors. Therefore, we applied the SILAC-controlled setup displayed in Figure 2B and used a stringent, sample-specific enrichment cut-off to call the interaction of a protein with DNA (Figure S3A, for details see Methods). This discovered 1805 candidate proteins, 1191 of which were present in three or more replicates (Figure S3B, Table S1). Figure 3A shows that 60 % carried a GO annotation for nucleic-acid binding (GO: DNA/RNA binding). To understand the contribution of specific protein classes to the DNA interactome, we used intensity-based absolute quantification (iBAQ), which estimates protein copy numbers in a sample from their combined peptide MS intensity (Schwanhäusser et al., 2011). As expected, the largest iBAQ contribution among proteins annotated as nucleic-acid binding came from histones, followed by proteins involved in mRNA processing and DNA repair (Figure S3C). Among proteins lacking an annotation for nucleic-acid binding the largest contribution came from lamins and other components of the nuclear lamina, followed by proteins involved in chromatin organization and cell division (Figure 3A & Figure S3D). Additionally, we identified a group of 278 proteins that could not be assigned to any of the common GO terms, representing a highly diverse set of molecular functions (Figure 3A). Overall, these were functions known to occur within chromatin, however, our DNA interactome highlighted specific proteins that came in direct contact with DNA when executing them. For example, we found proteins related to the ubiquitin-proteasome system such as PSM3, which forms an 11S regulatory complex (PA28γ) that promotes the degradation of TP53 and many other proteins involved in cell cycle progression, apoptosis and DNA repair (Thomas and Smith, 2022; Zhang and Zhang, 2008). Our data implied that PSM3 came in physical contact with DNA when targeting the proteasome within chromatin. Within the group of 278 proteins, 74 were linked to a genetic disease in OMIM, in total representing gene-disease associations for 99 genetic conditions (Amberger et al., 2009). Proteins with the most disease associations included the spectrin SPTAN1 and the protein phosphatase PTPN11 (aka. SHP2), which has been shown to associate with mitotic chromosomes during cytokinesis (Liu et al., 2016), in line with our observation that PTPN11 had direct access to DNA. Apart from these well-studied examples we found a number of uncharacterized proteins in this part of the DNA interactome, one of which was C5orf24. In a deep nuclear proteome from MCF7 cells (Figure S3E & Figure S3F, see Methods for details) C5orf24 ranked in the top 10 % of the most abundant proteins in the nucleus (Figure 3B). Remarkably, the entire sequence of C5orf24 constitutes a single domain of unknown function. DUF5568 is conserved across chordates and mostly found in single-domain proteins, yet, in rare cases in combination with a DEAD-box helicase domain (Paysan-Lafosse et al., 2023). Much of the C5orf24 sequence is predicted as disordered (88 % of amino acids with IUPRED score > 0.5) and AlphaFold shows very little tertiary structure for the human protein or its homologs in mice, fish or frogs (Figure S3G), suggesting that DUF5568 might either constitute an unknown, especially disordered ‘DNA-binding domain’ – in principle a DNA-binding disordered region (IDR) – or might participate in a phase-separated compartment that has direct access to DNA. When further comparing relative protein abundances between the MCF7 DNA interactome and the nuclear proteome, we noticed that some proteins ranked much higher than expected from their abundance in the nucleus (Figure 3B). For example, among the 117 transcription factors in our DNA interactome NACC1, CUX2, RARA, SOX4 and THAP10 had particularly good access to DNA in MCF7 cells while others such as TFAP2C, ZBTB7B or ZNF618 were excluded from it, confirming that transcription factor activity is highly regulated and often cannot be predicted from protein expression (Zaborowski and Walther, 2020). Overall, the direct DNA interactome provides a quantitative resource of proteins with physical access to DNA in MCF7 cells.

**Figure 3:**
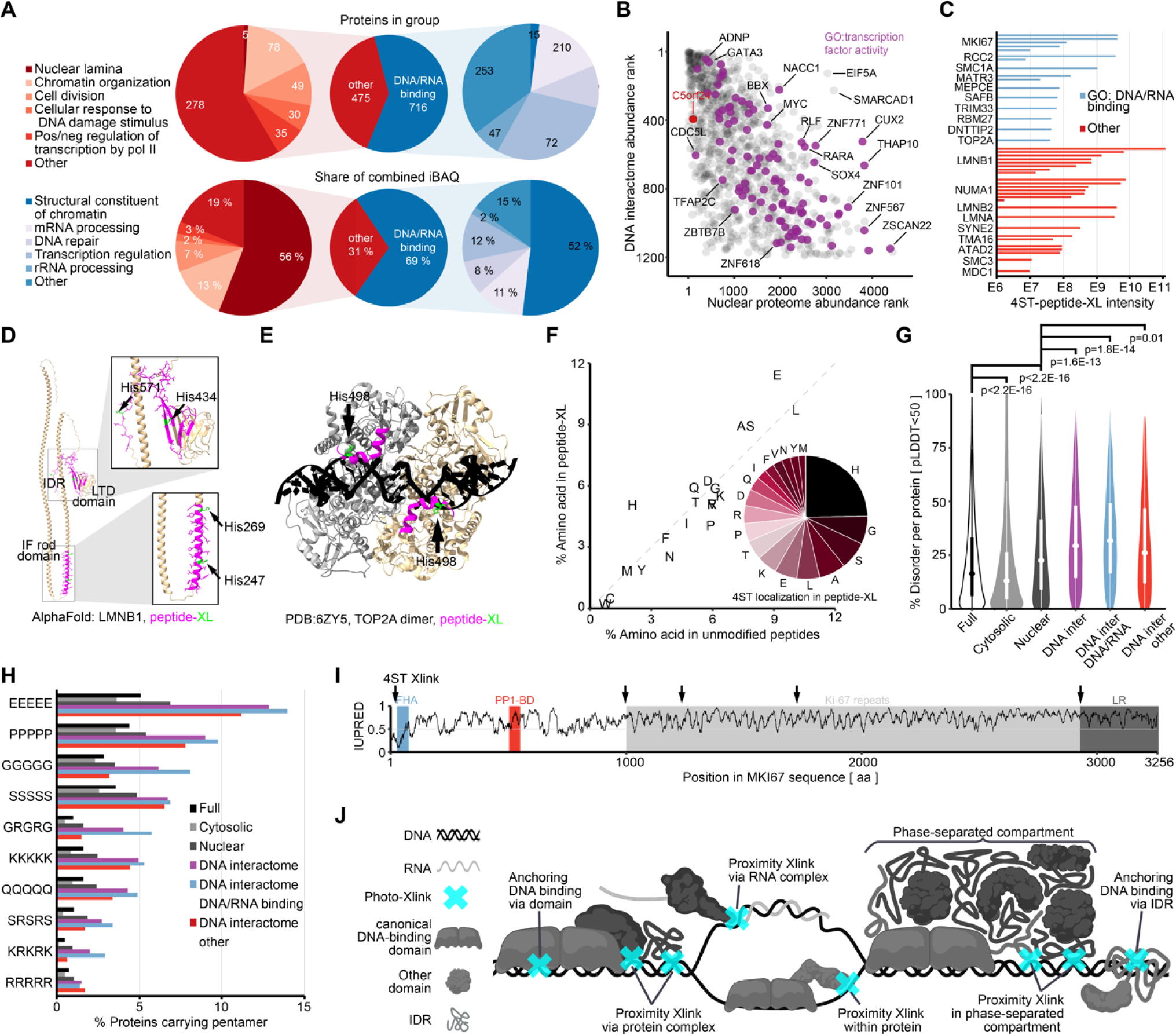
A census of proteins, domains and IDRs with direct access to DNA. A) Pie diagrams characterizing the constituents of the direct DNA interactome from MCF7 cells (see also Figure S3A&B). Proteins were first categorized according to their prior annotation for nucleic acid binding and subsequently assigned to the GO term with the largest combined iBAQ. Each protein contributes its iBAQ only once to the GO term it was first assigned to. B) Scatter plot comparing the relative abundance of proteins within the MCF7 nuclear proteome to their relative abundance in the DNA interactome. C) Bar plot comparing MS intensities of crosslinked DNA-peptide hybrids identified in the MCF7 DNA interactome. Displayed are peptides from the top ten proteins with the highest intensity crosslinks with and without prior GO annotation for nucleic acid binding. D) AlphaFold structure prediction for LMNB1 displaying DNA crosslinks from C. Identified 4ST-crosslinked peptides are displayed in magenta and the modified amino acid in green. E) Cryo-electron microscopy structure of the TOP2A DNA-binding/cleavage domain dimer in complex with its DNA (black) substrate (PDB 6ZY5). The identified 4ST-crosslinked peptide is indicated in magenta, histidine crosslinking site in green. F) Scatter plot comparing amino acid frequencies in unmodified peptides of the MCF7 DNA interactome to nucleotide-crosslinked peptides found in the same data. The inserted pie diagram indicates the relative occurrence of amino acids at the crosslinking site as reported by MSfragger. G) Violin plots comparing the percentages of amino acid positions predicted as disordered by AlphaFold (pLLDT<50) (Akdel et al., 2022). Testing occurred with a two-sided Kolmogorov-Smirnov test. Proteins in the MCF7 full proteome (full) are compared to the cytosolic and nuclear proteome (see Figure S3D), the DNA interactome (DNA inter) and its parts annotated as nucleic-binding (DNA/RNA) or lacking this annotation (no DNA/RNA). H) Barplot comparing the occurrence of the most abundant disordered pentamers between the same groups as in G. See also Figure S4L. I) Lineplot showing IUPRED scores of MKI67 along its entire sequence. DNA crosslinking sites identified in the MCF7 DNA interactome are indicated by arrows (4ST Xlink). J) Schematic summary of protein-DNA interactions observed among proteins with direct access to DNA. For details see text.

### Structural properties of proteins in direct contact with DNA

Chromatin has been shown to be solid-like in living cells, providing a nucleosome scaffold for protein and RNA that creates an elastic gel (Strickfaden et al., 2020). The term ‘DNA binding’ in the conventional sense describes a physical protein-DNA interaction with the purpose of anchoring a protein to the nucleic acid sequence, e.g. when a PHD zinc-finger domain binds DNA to physically anchor a transcription factor to a gene promotor and elicit transcription. However, due to the gel-like nature of chromatin, proteins can also interact with DNA circumstantially and without ‘binding’ to it, simply because they come into DNA proximity as part of a complex or a phase-separated compartment (Boija et al., 2018; Sabari et al., 2018). We were interested to see if there were common sequence features among proteins in the DNA interactome that might grant them access to the genome and anchor them there. When comparing the occurrence of domains in the MCF7 DNA interactome to their occurrence in the MCF7 full proteome, we found strong enrichment for canonical DNA and histone-binding domains, such as the PHD-type and C2H2-type zinc-finger, the SAP domain, DEAD/DEAH box helicase, as well as the chromodomain, bromodomain and the SANT domain (Figure S3H). Interestingly, the RNA recognition motif (RRM) domain was also strongly enriched in the DNA interactome. The RRM has been shown to bind DNA as well as RNA *in vitro*, albeit, only in their single-stranded form (Chen et al., 2021). For direct evidence of protein-DNA crosslinking in RRM domains we searched the DNA interactome for 4ST-crosslinked peptides and indeed found crosslinking sites mapping to the RRM domains of MATR3, TRA2B, SRSF2, SRSF11 and RNPS1, implying that the RRM can interact with DNA in living cells. Overall, 162 4ST-crosslinked peptides from 83 proteins could be identified in the MCF7 DNA interactome (Table S2). The protein with the most nucleotide-crosslinks identified, and at the same time contributing the 4ST-modified peptides with the highest MS intensities, was LMNB1 (Figure 3C & Figure 3D). We located a 4ST-modified peptide in the DNA-binding domain of the type-II topoisomerase TOP2A, whose crystal structure complexed with a DNA substrate in Figure 3E illustrates the histidine the crosslink could be localized to. In fact, all 4ST modifications in LMNB1 were localized to histidines (Figure 3D), too, and so were 25 % of all other identified crosslinks followed by glycine, serine and alanine (Figure 3F). Only 61 out of the 162 4ST-modified peptides could be mapped to annotated domains whereas many localized to IDRs (Table S2). This has been observed in crosslinking studies investigating protein-RNA interactions before (Bae et al., 2020; Castello et al., 2016; Kramer et al., 2014; Trendel et al., 2019). In order to understand what role protein disorder played in the DNA interactome we used the IUPRED score (Erdos et al., 2021) and AlphaFold’s confidence score pLDDT (Akdel et al., 2022) as proxies for disorder. Both scores were in good agreement that proteins in our deep nuclear MCF7 reference proteome were significantly more disordered than the entire MCF7 full proteome (two-sided Kolmogorov-Smirnov test, p<2.2E-16), and accordingly, proteins in the deep cytosolic proteome significantly more folded (p<2.2E-16, Figure 3G & Figure S3I). Compared to the nuclear proteome, proteins in the DNA interactome were even more disordered (p=1.6E-13) and proteins within the DNA interactome carrying an annotation for nucleic-acid binding showed the strongest disorder (p=1.8E-13). In median, the combined length of disordered positions in a cytosolic protein amounted to 53 amino acids, in a nuclear protein to 115, and in a protein contained in our DNA interactome to 184 (AlphaFold pLDDT<50). Overall these findings implied that proteins able to access DNA were often much more disordered than proteins in their surrounding. We emphasize, however, that protein disorder was not at all predictive for the subcellular localization of a protein to the nucleus or direct DNA interaction. We asked if there were commonalities among the disordered features in proteins accessing DNA and how the amino acid sequence of these proteins might be different to the rest of the proteome. Interestingly, the amino acid composition of proteins in the DNA interactome was virtually identical to the composition of proteins in the MCF7 full proteome (Figure S3J). We looked for IDRs directly and counted the occurrence of pentamer sequences containing any permutation of the amino acids G, S, D, Q, P, E, K, and R, typically associate with protein disorder (Van Der Lee et al., 2014). Indeed, this discovered homopolymeric sequences of E, P, S, G, K and Q, as well as RS and RG-repeats strongly enriched in the DNA interactome compared to the MCF7 full proteome (Figure S3K). The occurrence of these repeats again tracked with the proximity of proteins to DNA (Figure 3H). In combination the ten repeat sequences displayed in Figure 3H occurred in 36 % of proteins in our DNA interactome and only 13 % of the MCF7 cytosolic proteome. By far the most frequent homorepeat in our DNA interactome were stretches of poly-E, whose occurrence in proteins involved with chromatin organization and transcription has been recognized before and is conserved from yeast to human (Chavali et al., 2020; Lee et al., 2022; Soto et al., 2022). Among disordered motifs unique to a single protein the Ki-67 repeats of the highly disordered MKI67 (85 % of amino acids with IUPRED score > 0.5) stood out, to which several 4ST-peptide crosslinks with very high MS intensity could be mapped (Figure 3I & Figure 3C). Figure 3J summarizes the protein-DNA interfaces implied by our DNA interactome and the nucleotide-crosslinked peptides it contained. First, many conventional DNA interactions were represented in the DNA interactome including over one hundred transcription factors carrying canonical, globular DNA-binding domains. Second, we observed numerous IDR interactions with DNA, some of which might mitigate anchoring ‘DNA binding’ and some of which might occur circumstantial. This is best exemplified on MKI67, whose disordered C-terminal LR domain binds DNA *in vitro* and has been shown to be necessary and sufficient for anchoring the protein to chromosomes in cells (Sun and Kaufman, 2018). Despite crosslinks in the C-terminal IDR we found crosslinks in the disordered core Ki-67 repeat regions and even the N-terminal FHA domain (Figure 3I, Table S2), implying that all these parts had access to DNA while not anchoring to it. During mitosis MKI67 engulfs chromosomes as a surfactant that prevents their mutual collapse (Cuylen et al., 2016), illustrating how circumstantial DNA crosslinking might occur when parts of a protein are permanently kept in proximity to DNA within phase-separated compartments. Finally, we observed a number of interactions that were best explained by circumstantial crosslinking of proteins being brought into proximity of DNA via interaction with other proteins or RNA, e.g. bromodomain proteins interacting with histones or splicing factors with nascent mRNA.

### Quantification of transcription factor-binding to DNA during cellular perturbations

Since the signaling activity of transcription factors requires their interaction with the genome, observing changes in their DNA engagement could in turn indicate an involvement in the genomic regulation towards a specific perturbation. For instance, it is known that the estrogen receptor alpha (ESR1) resides in the cytosol until it encounters its ligand estrogen, triggering translocation into the nucleus where ESR1 binds specific regions in the genome and activates a specific transcriptional program (Hah et al., 2011). We were interested to see if by quantitatively comparing DNA interactomes between untreated and estrogen-treated cells, we could recapitulate this genomic regulation. Therefore, serum-starved MCF7 cells were exposed to a high dose (10 nM) of estrogen (17β-estradiol) for 45 minutes and their DNA interactome compared to mock-treated control cells (Figure 4A). Indeed, we observed a 30-fold increase of ESR1 in the DNA interactome after estrogen stimulation (Figure 4B, Table S3), which highlighted it as the most significantly changed protein in the differential analysis (adjusted p=2.6E-42, negative binomial model (NBM) testing, see Methods for details). In total, around 9% of the 1814 quantified protein-DNA interactions showed significant changes exceeding 2-fold (adjusted p<0.01). We observed increased DNA binding of known ESR1 interactors such as RNA polymerase II (POLR2A, adj. p=2.1E-6, Figure 4B) or the chromatin remodeler SMARCA4 (aka. BRG1, adj. p=2.7E-5), a member of the SWI/SNF complex that has been shown to be recruited by ESR1 to estrogen-responsive genes and to be required for effective transcription activation (DiRenzo et al., 2000; Vaart et al., 2020). Using iBAQ as approximation for protein copy numbers on DNA, we found the repressors SET, CBX3 and CBX5 showing the largest absolute loss in interaction with DNA (Figure 4C, Table S3). Bound to H3K9 methylation CBX3 and CBX5 promote heterochromatin formation (Zeng et al., 2010), whereas SET represses histone acetylation (Seo et al., 2001), suggesting that their estrogen-induced release from DNA prepared decondensation of chromatin and activated transcription in the serum-starved cells. The most substantial absolute increase in protein abundance on DNA was observed for the nucleolar transcription factor UBTF, a crucial regulator of rRNA synthesis (Jantzen et al., 1990), alongside other proteins implicated in ribosome biogenesis, such as SUMO2 and SUMO3, suggesting an immediate estrogen-induced stimulation of ribosome and protein production.

**Figure 4:**
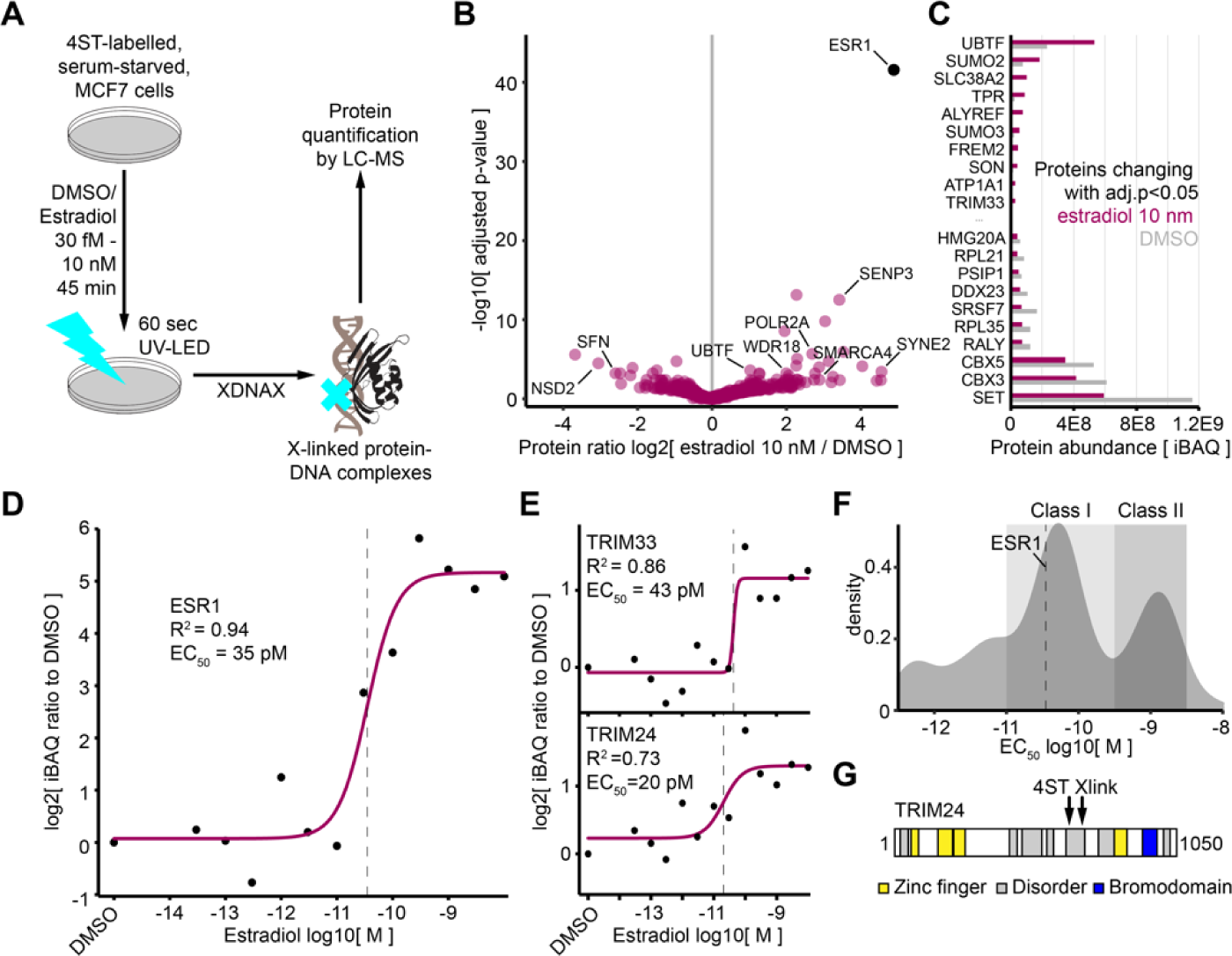
Quantitative comparisons of DNA interactomes from breast cancer cells treated with estrogen. A) Experimental outline for the comparison of proteins interacting with DNA before and after 45 minutes of estrogen treatment in 48-hour serum-starved MCF7 cells. B) Volcano plot illustrating changes in the DNA interactome after 45 minutes of high-dose estrogen exposure (10 nM 17β-estradiol). C) Bar plot comparing the most extreme absolute changes in the abundance of proteins interacting with DNA in the same experiment as B. D) Exemplary dose-response curve derived from the relative protein abundance of the estrogen receptor ESR1 in the DNA interactome of MCF7 cells at increasing estrogen concentrations. E) Same as in D but for the proteins TRIM33 and TRIM24. F) Density plot of half-effective concentrations for the association of proteins with DNA during estrogen exposure (see Figure 5D). Two classes of proteins are identified changing their interaction at low (class I) or high (class II) estrogen concentration. G) Domain structure of TRIM24 indicating the estrogen-induced DNA crosslinks in its core IDR.

In reference to our recently reported methodology for the deconvolution of drug action via dose-dependent quantification of protein modifications (decryptM, (Zecha et al., 2023)) or via full proteomes (decryptE, Berner et al., in press), we expanded the measurements and treated MCF7 cells with eleven additional concentrations of estrogen ranging from 10 nM to 30 fM for 45 minutes before quantifying their photo-crosslinked DNA interactome (Figure 4A). Using a log-logistic dose-response model, we were able to derive half-effective concentrations (EC_50_) for the engagement of 1483 proteins with DNA (Figure S4A, Table S3). The best fit in the relevant concentration range was for ESR1 (R^2^=0.94, Figure 4D), showing an EC_50_ of 35 pM for its engagement with DNA, which is close to the K_d_ of 50 pM reported from a cell-free radioligand assay (Kuiper et al., 1998). We observed numerous proteins changing their interaction with DNA at similar or slightly higher effective concentrations than ESR1 at around 100 pM estrogen, such as the chromatin organizers TRIM33 and TRIM24, which doubled their abundance on DNA in this estrogen concentration range (Figure 4E). TRIM24 has been shown to be a co-activator of ESR1 during estrogen stimulation of MCF7 cells, and its overexpression has been correlated with poor breast (Tsai et al., 2010) and prostate cancer outcomes (Groner et al., 2016). Other changes in protein-DNA interactions within a similar concentration range as ESR1 included increased interaction of the transcription activators SMARCA4, UBTF, and BPTF, as well as decreasing interaction of the repressor CDX3 (Figure S4B). In addition, we found a second group of proteins with approximately ten times higher effective concentrations (Figure 4F, class II). These included genes known to be transactivated by ESR1, implying that estrogen doses higher than the effective concentration of ESR1 had increased their expression, leading to increased binding of their protein product to DNA. One interesting example for this was SYNE2, which showed increased binding to DNA only at an EC_50_ of 1.9 nM, yet, a robust 23-fold increase at 10 nM concentration (adj. p=3.8E-4, Figure 4B & Figure S4C). In humans, the SYNE2 gene is located on chromosome 14 immediately next to the ESR2 gene (estrogen receptor β, Figure S4D), whereas at the paralogue locus on chromosome 6 the SYNE1 gene is located immediately next to the ESR1 gene (estrogen receptor α, Figure S4E). ENCODE Chromatin immunoprecipitation followed by sequencing (CHIP-Seq) data for ESR1 in breast cancer cells exposed to 10 nM estrogen for one hour shows strong recruitment of ESR1 to the ESR2/SYNE2 locus (Dunham et al., 2012), suggesting potential crosstalk between the genes and their protein products. In our SILAC-controlled DNA interactome SYNE2 was confidently identified as DNA interacting in all replicates (Table S1) and contributed one of the 4ST-crosslinked peptides with the highest intensities, further corroborating that its interaction with DNA can be direct (Figure 3C). Across all samples in our estrogen series we identified 346 4ST-crosslinked peptides from 255 proteins (Table S2), some of which recapitulated the estrogen-induced changes we had observed earlier. For example, we identified two DNA-crosslinking sites in a central IDR of TRIM24 that were only detected after exposure to estrogen (Figure 4G), confirming its recruitment and direct interaction with DNA (Figure 4E). Overall the quantitative comparison of DNA interactomes during estrogen treatment validated our approach for the unbiased elucidation of genomic regulation during cellular perturbations on a time scale of minutes.

### Genomic regulation in response to three clinically relevant genotoxic drugs

In addition to understanding how the genome is read and regulated by interactions with proteins, there is great interest in understanding how these interactions organize its maintenance and repair. This is particularly relevant in respect to chemotherapeutic cancer treatments, where genotoxic drugs form an important pillar of clinical pharmacology aiming to halt the growth of fast dividing cells by damaging their DNA (Swift and Golsteyn, 2014). We considered three commonly applied genotoxic compounds – etoposide, cisplatin and oxalipatin – which are well-characterized and known to act by distinct modes of action (Bruno et al., 2017; Swift and Golsteyn, 2014). Live cell imaging on MCF7 cells indicated that all drugs reached their maximal potential to halt proliferation and induce apoptosis 24 hours after addition (Figure S5A), indicating that abundant DNA damage had been produced at this point. We derived DNA interactomes from cells after 24 hours of treatment with each drug and quantified them against mock-treated control cells (Figure S5B), comparing approximately 2500 proteins per condition (Table S4). Although the majority of protein-DNA interactions remained constant, all three genotoxic compounds resulted in highly significant and specific interaction changes (Figure 5A-C). Common to all treatments were strongly increased DNA interactions of key cell cycle regulators such as CDKN1A (aka. p21, Figure S5C), and DNA repair proteins such as TP53BP1, SIRT7 and in particular HINT1, which helps to resolve γ-H2AX phosphorylation at DNA damage sites after repair (Li et al., 2008). Recruitment of these proteins to DNA implied that DNA damage had occurred and triggered cell cycle arrest. The most significant loss in DNA interaction across all treatments was observed for a component of the telomerase complex, NHP2, mutations of which can cause premature ageing in humans (Astorgues-Xerri et al., 2014). NHP2 was only detected on DNA in untreated cells, indicating that any genotoxic drug treatment led to decreased telomerase activity and contributed to damage-induced senescence (Zhang et al., 2002). All drugs also shared the recruitment of proteasome components like PSMC2, which was absent from DNA in untreated cells. The proteasome has been shown vitally involved in the repair of drug-induced DNA-protein crosslinks (Essawy et al., 2023), and PSMC2 forms part of the 19S regulatory particle that recognizes target proteins to funnel them into the 20S catalytic core (Kamber Kaya and Radhakrishnan, 2021). Under cisplatin treatment PSMC2 showed the most significant abundance increase in the entire experiment (adj. p=1.4E-108, NBM testing, Figure 5B) and was more than five times more abundant on DNA than in the other drug treatments (Figure S5C), underscoring the extensive DNA-protein crosslinking occurring under the drug (Stingele et al., 2017).

**Figure 5:**
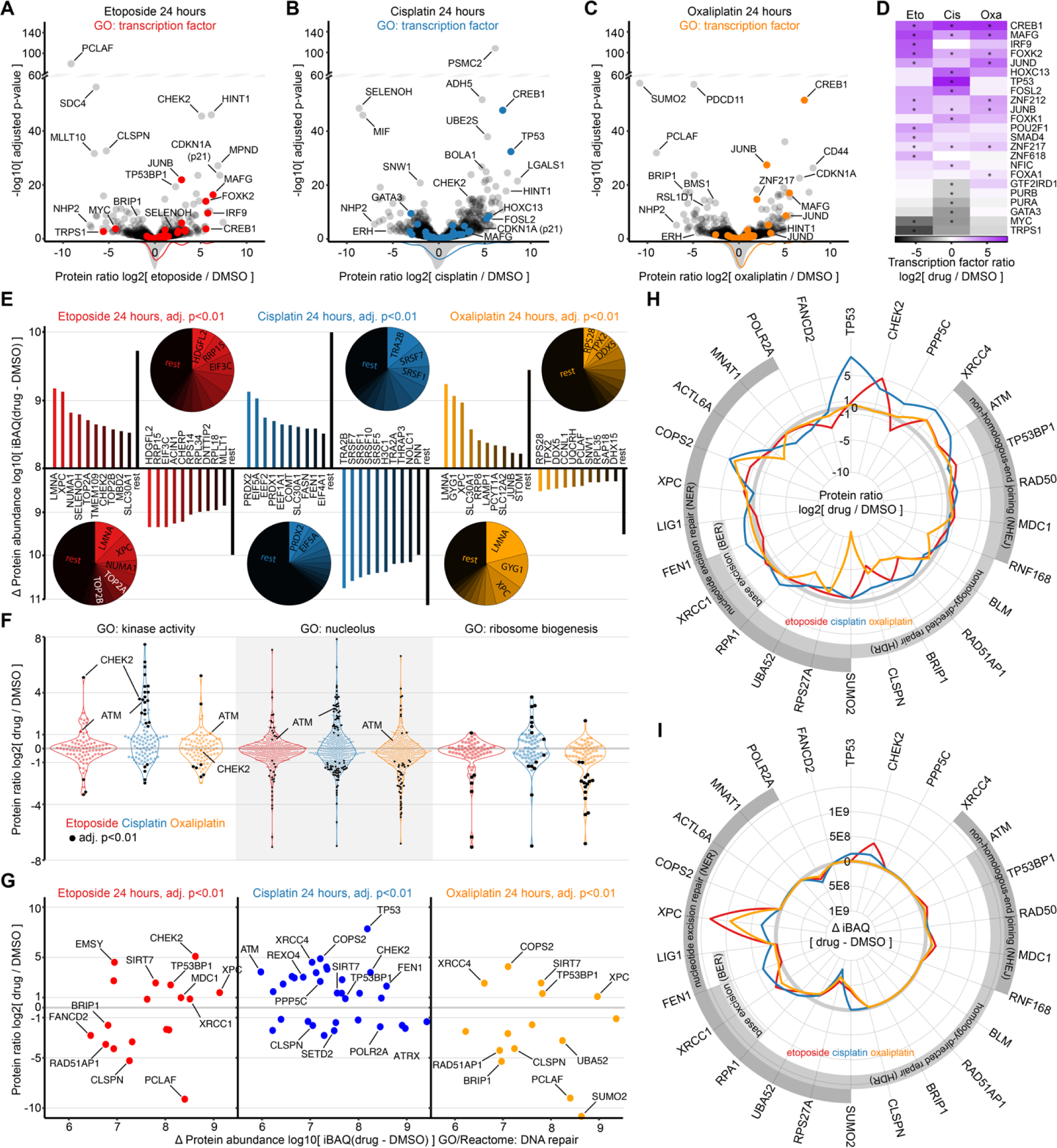
Changes in the direct DNA interactomes from breast cancer cells exposed to different genotoxic drugs. A)-C) Volcano plots showing differences in the DNA interactomes of 24-hour mock-treated MCF7 cells (DMSO) to cells treated with etoposide, cisplatin or oxaliplatin at 100 µM concentration. Missing values were imputed to include proteins without DNA interaction before or after treatment (e.g. TP53, CREB1, PCLAF etc., see Table S4). D) Heatmap comparing foldchanges of transcription factor-DNA interactions between treatments. All transcription factors with significantly changed interaction in at least one treatment are displayed (adj. p<0.01 indicated by *). E) Barplots showing the top ten absolute increases or decreases in protein abundance within the indicated DNA interactome. Compared are only proteins whose relative change was also found significant (adj. p<0.01). Pie diagrams illustrate the relative contribution of each protein. F) Beeplot comparing changes in the DNA interactomes for selected proteins carrying GO annotations for kinase activity, nucleolar localization or ribosome biogenesis. G) Scatter plot relating absolute and relative abundance changes of DNA repair proteins in the DNA interactome of cells treated with the indicated drug. Within each treatment only DNA repair proteins with significantly changed interactions are shown (adj. p<0.01). H) Radar plot comparing relative abundance changes in the DNA interactome for proteins participating in specific repair pathways (Reactome annotation). Only proteins with significantly changed interactions under any of the three drugs are displayed (adj. p<0.01). I) Same as in H but comparing absolute protein abundance changes between the DNA interactomes.

In comparison to all changes in the DNA interactomes, we observed a significant increase in transcription factor binding for each of the drugs (p ≤ 0.01 for all three treatments, two-sided Kolmogorov-Smirnov test), implying intense genomic regulation in response to genotoxic damage. Figure 5D shows that among the 23 transcription factors exhibiting significant changes in any of the treatments (adj. p<0.01, NBM testing), 17 increased binding whereas only six decreased it. Strongest decrease in DNA interaction for a transcription factor was observed for MYC, whose expression is known to be reduced upon genotoxic stress in MCF7 and other cell lines (Castell et al., 2022). Under cisplatin we also observed strongly reduced DNA interaction of GATA3, which is used as biomarker for metastatic potential in clinical staging of breast cancer biopsies (Adomas et al., 2014). We found four transcription factors with significantly increased DNA interaction across all treatments (adj. p<0.01), including the stress mediators MAFG and CREB1, which were undetected on DNA under mock treatment and whose knockdown reportedly confers resistance to cisplatin (Kim et al., 2019; Shi et al., 2004). In addition, drug-specific increases in DNA binding were observed for JUND, JUNB and FOSL2, all of which share the ability to dimerize into various manifestations of the transcription activating complex AP-1 via cross-family dimerization (Hai and Curran, 1991). Another notable drug-specific change of transcription factor interactions with DNA was a significant increase in IRF9 in response to etoposide (p=1.3E-14), which is a known inducer of type-I interferon expression, autoinflammation, and resistance to DNA damage (Cheon et al., 2013), as well as a highly significant rise in TP53 binding to DNA in response to cisplatin (p=2.1E-38). In mock-treated cells, TP53 was undetectable on DNA, ranked in the bottom 30% of protein abundances under etoposide or oxaliplatin treatment, but in the top 20% of protein abundances under cisplatin stress (Table S4). This increase was only surpassed by JUNB in oxaliplatin-treated cells, whose gain on DNA ranked in the top ten of all abundance increases (Figure 5E). When comparing absolute changes in protein abundances between the DNA interactomes, we observed large amounts of peroxiredoxins accumulating on DNA specifically in cisplatin-treated cells, where PRDX1 and PRDX2 combined made up for more than 10 % of the entire gain in iBAQ compared to the mock-treated control (Figure 5E). Peroxiredoxins protect cells from oxidative stress and have been shown to offer cisplatin resistance (Yoo et al., 2002). PRDX2 interaction with chromatin has been reported before, where it was preferably observed in a multimeric conformation. Intriguingly, under conditions of oxidative stress PRDX2 forms a decameric toroid (Pastor-Flores et al., 2020; Somyajit et al., 2017), which by proportion would be large enough to accommodate the DNA double helix (Figure S5D). In summary, genotoxic stress changed the DNA interactome of MCF7 cells profoundly, creating a characteristic landscape of interactions with transcription regulators, metabolic enzymes, repair and structural proteins specific to each drug.

### Mining damaged DNA interactomes for repair mechanisms and potential vulnerabilities

The three genotoxic drugs we applied have long been among first-line treatments for various cancer indications in the clinic (Swift and Golsteyn, 2014). We were interested whether our DNA interactomes could recapitulate repair pathways counteracting the drug-induced DNA damage, and if so, whether this suggested cancer vulnerabilities that might be exploited in potential combination treatments. Interestingly, principle component analysis (PCA) indicated that protein abundances in the DNA interactomes from cells treated with etoposide or oxaliplatin were more similar than from cells treated with cisplatin (Figure S5E). However, the recruitment of individual DNA damage response proteins was quite distinct between treatments. For instance, the central damage-relaying kinases ATM and CHEK2 were strongly recruited to DNA in response to cisplatin, only CHEK2 in response etoposide, and neither in response oxaliplatin (Figure 5F). For a consistent comparison between the three drugs, we focused on the best-populated DNA repair pathways in the DNA interactomes and compared them directly (Figure 5G-I). Etoposide is known to stabilize a usually transient covalent interaction in the enzymes TOP2A and TOP2B during their catalytic cycle of DNA cleavage and re-ligation to release superhelical tension, resulting in single and double-strand breaks (Muslimović et al., 2009). In line with previous reports on the repair of etoposide-induced damage (Biard, 2007; Olivieri et al., 2020), a direct comparison of DNA damage proteins revealed signatures for the repair of DNA double-strand breaks via the non-homologous end joining (NHEJ) pathway, as well as repair of base damage via the base-excision repair (BER) and nucleotide-excision repair (NER) pathways (Figure 5H). Several proteins specifically involved in homology-directed repair (HDR), such as CLSPN, BRIP1 and RAD51AP1 were depleted from DNA under etoposide treatment, indicating that this pathway was suppressed. This was corroborated by strong recruitment of the HDR inhibitor EMSY to DNA (adj. p=1.1E-19, NBM testing, Figure 5G) (Marzio et al., 2022) alongside depletion of the HDR promotor HDGFL2, which showed the largest absolute decrease in protein abundance on DNA of all proteins after etoposide treatment (Figure 5E)(Baude et al., 2015). Active removal of topoisomerases trapped on DNA has been shown to occur in MCF7 cells within a few hours after addition of etoposide and is triggered by the SUMO-ligase PIAS4, followed by ubiquitination and proteasomal degradation (Sun et al., 2020). After 24-hour etoposide stress no increased interaction of PIAS4 with DNA was observed, whereas TOP2A and TOP2B were among the ten proteins with the largest absolute gains in protein abundance (Figure 5E), representing 2-fold increases over untreated cells (adj. p<0.006, Table S4). We derived additional DNA interactomes from cells treated for four hours with etoposide that showed 6-fold TOP2A and TOP2B accumulation (Figure S5F), strong TOP2A phosphorylation (Figure S5G), as well as significant recruitment of PIAS4 (adj. p<0.002, Figure S5H). We concluded that active removal of trapped topoisomerases was especially strong while cells were still proliferating and DNA synthesis produced superhelical tension, requiring strong type-II topoisomerases activity (Figure S5A).

Short term oxaliplatin treatment in the range of several hours has been shown to stop rRNA synthesis and cause ribosome biogenesis stress (Bruno et al., 2017), whereas longer treatment of up to one day causes DNA damage and disintegration of nucleoli (Schmidt et al., 2022; Sutton and DeRose, 2021). Indeed, after 24 hours of oxaliplatin we observed an extensive loss of nucleolar proteins from the DNA interactome (p= 6.4E-09, two-sided Kolmogorov-Smirnov test), including many proteins involved in ribosome biogenesis (p=4.3E-04, Figure 5F). In oxaliplatin as well as etoposide-treated cells the damage sensor XPC ranked among the top three absolute iBAQ increases (Figure 5E), whereas only a slight but insignificant increase of XPC was observed under cisplatin (Figure 5I). Just like etoposide, oxaliplatin led to strong depletion of the HDR-specific proteins CLSPN, BRIP1 and RAD51AP1 from DNA, again indicating that this pathway was avoided (Figure 5G & Figure 5H).

Cisplatin led to strongly increased DNA interaction of the NHEJ marker XRCC4, the BER/NER proteins RPA1, FEN1 and LIG1, as well as the key DNA damage relaying kinase ATM (Figure 5G & Figure 5H), overall confirming its ability to exert the most severe damage among all three drugs (Figure S5A). Apart from ATM and CHEK2 various other kinases accumulated on DNA under cisplatin treatment (Figure 5F), including RPS6KB1 (adj. p=2.6E-13), AKT1 (adj. p=0.008) and PRKCD (adj. p=0.009), all of which have been shown to be involved in cell cycle control or the DNA damage response. Notably, CHEK2 showed one of the largest absolute increases in protein abundance for both cisplatin and etoposide, while completely unchanged under oxaliplatin (Figure 5I). We used the DNA interactome of cisplatin-treated cells to identify potential sensitivity factors for combination treatments. The DNA repair enzyme showing the largest absolute increase in protein abundance on DNA was FEN1 (Figure 5I & Figure S5I), a 5’ flap endonuclease central to long-patch base-excision repair, which has in fact been associated with cisplatin resistance before (He et al., 2017; Ward et al., 2017). Two other enzymes reportedly involved in DNA repair were specifically enriched under cisplatin stress, the phosphatase PPP5C and the exonuclease REXO4 (Figure S5I). We used siRNAs to knock down each protein in MCF7 cells and followed their proliferation in the presence of increasing cisplatin concentrations by live cell imaging (Figure S6A). FEN1 and PPP5C knockdown strongly increased the induction of apoptosis (Figure 6A & Figure 6C) and cell death (Figure 6B & Figure 6D) at the cisplatin concentration used to derive our DNA interactome (100 µM), whereas REXO4 depletion led to de-sensitization at higher cisplatin concentrations (Figure S6B & Figure S6C). This implied that FEN1 and PPP5C but not REXO4 could represent potential drug target candidates alongside cisplatin. Overall, these findings confirmed that DNA interactomes can be leveraged to compare DNA-damaging drugs mechanistically and identify weaknesses in the DNA repair machinery of cancer cells.

**Figure 6:**
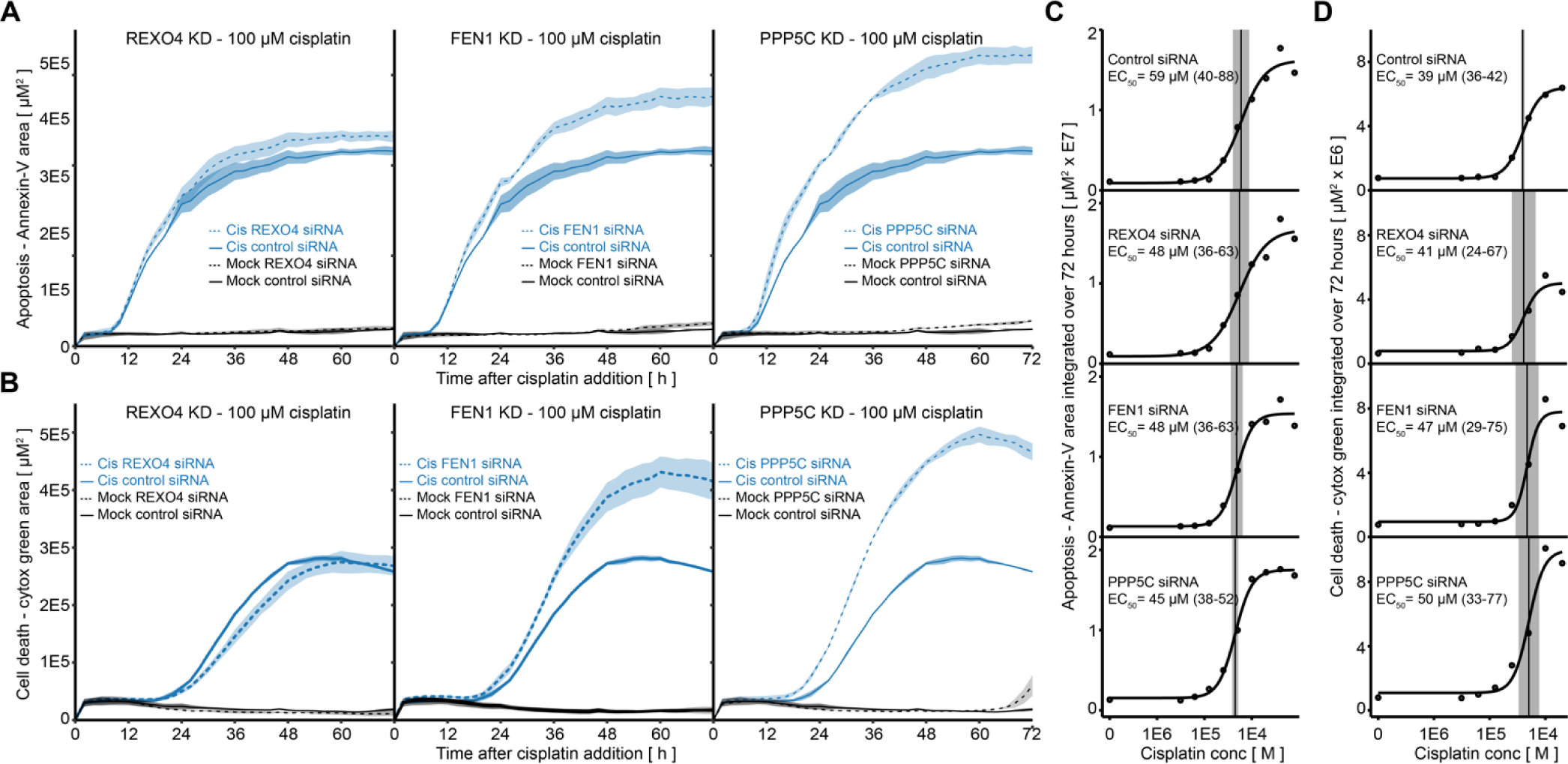
Live cell imaging time series evaluating DNA interactors as cisplatin sensitivity factors. A) Lineplots following the induction of apoptosis in MCF7 cells with knockdowns for REXO4, FEN1 or PPP5C using fluorescence-labelled Annexin-V. One uniform pool of cells was transfected with siRNAs 24 hours before addition of 100 µM cisplatin (cis) or drug-free media (mock). Bold lines indicate cells transfected with control siRNAs, dotted line for one specific siRNA target, shading one standard deviation. B) Same as in A but for the quantification of dying cells using the fluorescence dye cytotox green. C) & D) Dose-response curves using the integrated florescence signal over 72 hours to determine half-effective concentrations for cisplatin to induce apoptosis (C) or cell death (D) during the duration of the indicated knockdown in MCF7 cells.

## Discussion

### A photo-crosslinking system for the study of direct protein-DNA interactions in living cells

Investigating proteins accessing the genome to understand how the stored information is read and put into practice has been a long-standing quest. For example, the ENCODE project has mobilized massive efforts to describe the interaction of hundreds of DNA-binding proteins with the human and mouse genome in numerous cell lines, tissues and developmental stages (Abascal et al., 2020a). Chemical crosslinking of protein and DNA with formaldehyde (FA) has played a major role in this achievement, contributing to almost half of the over 8000 sequencing assays reported by ENCODE in 2020 (Abascal et al., 2020b). The high protein content of FA-crosslinked chromatin has posed challenges for its purification for proteomic analysis, as classic biochemical isolation procedures are very effective at purifying protein or nucleic acids alone but not a crosslinked hybrid molecule sharing physicochemical features of both. During metabolic labeling with the photo-activatable 4ST only about 0.1 % of thymidine bases are replaced (Reelfs et al., 2011), implying that large parts of the DNA molecule remain non-crosslinked to protein after irradiation. We capitalized on this sparse crosslinking by using a of a series of standard DNA isolation procedures during XDNAX that allowed for very stringent removal of non-crosslinked protein from DNA, resulting in extracts of protein-crosslinked DNA with exceptionally high purity. Obvious drawbacks of our photo-crosslinking approach include the requirement for cellular proliferation to incorporate 4ST into the genome via DNA synthesis, as well as intense photo-activation with UV light.

With the introduction of the LED-based irradiation system UVEN, we reduced the time for protein-DNA crosslinking in living cells to one minute at four degrees Celsius, preventing cellular reactions that might change the composition of the DNA interactome in the crosslinking process. This is markedly different to FA crosslinking, which requires at least ten minutes of fixation at room temperature. To achieve similar crosslinking with a conventional 365 nm bulb-based device would require at least 100 times longer irradiation time. This means cells would need to remain outside normal culture conditions and under irradiation for almost two hours, what is incompatible with the survival of most cultured cells and would likely influence the outcome of the crosslinking experiment. The UVEN irradiation system is being developed as an open science project, for which we provide detailed plans, construction manuals and software under the Creative Commons license at wwww.uven.org, with the aim of making photo-activation in living cells accessible to a broad research community. Several UVEN devices have been constructed and are currently tested by different laboratories for photo-crosslinking studies of protein-DNA, protein-RNA, protein-drug as well as protein-protein interactions for proteomic, genomic and transcriptomic applications, showcased in an upcoming publication.

### Physical access to the genome from a proteomic perspective

Chromatin accessibility is determined by the topology of nucleosomes and the association of chromatin-binding proteins and RNA around them (Klemm et al., 2019). Accessibility has primarily been investigated from a DNA-centric perspective, where sequencing techniques such as DNase I hypersensitive site sequencing (DNase-Seq) or Assay for transposase-accessible chromatin using sequencing (ATAC-Seq) have been applied to assess what loci in the genome are open for protein binding. Various proteomic approaches have explored the overall composition of chromatin, however, what proteins within the chromatin gel physically interact with DNA has so far remained unclear (Ginno et al., 2018; Lewis and Laemmli, 1982; Van Mierlo and Vermeulen, 2021; Nakamura et al., 2021; Ohta et al., 2010; Rafiee et al., 2021). Using photo-crosslinking, we derived an atlas of human proteins in direct contact with DNA that offered insights on how proteins access the genome. The majority of proteins in this DNA interactome carried canonical DNA, RNA or histone-binding domains, underscoring the importance of globular domains for DNA access by permanently anchoring proteins to the DNA molecule directly or indirectly. In addition, we found that proteins with direct DNA access often carried large IDRs. Specifically for transcription factors it has been recognized before that many of them comprise large IDRs, which have been shown to modulate the DNA sequence they bind to or their ability to activate transcription (Ahn et al., 2021; Boija et al., 2018; Brodsky et al., 2020; Xing et al., 2023). While transcription factors typically feature IDRs alongside globular DNA-binding domains, we observed various proteins in direct contact with DNA that were mostly disordered and did not carry any globular DNA-binding domains at all. Thus, for future classification we found it useful to differentiate between DNA-binding IDRs that provide DNA anchorage, IDRs that allow proteins to participate in phase-separated compartments adjacent to DNA, and any third type of IDR that serves another function (Figure 3J). In our DNA interactome disordered proteins were enriched in comparison to the surrounding nuclear proteome, suggesting that some IDRs might facilitate direct access of proteins to DNA. We hypothesize that in the competition for a direct interface with the DNA molecule, proteins with certain IDRs have an advantage: They can participate in specific phase-separate compartments that already exist around DNA to gain access and, subsequently, use their DNA-binding or histone-binding domains to anchor and stay in place (Brodsky et al., 2020; Xing et al., 2023). Conversely, once a protein has bound its target DNA locus with a dedicated DNA-binding domain, its IDRs can establish a phase-separated compartment that facilitates access of specific auxiliary proteins in its vicinity (Boija et al., 2018).

Future studies investigating the DNA interactome of other human cell lines and other species will provide a more detailed picture about the variability of DNA access and how it evolved. The recent discovery of histone-like proteins in bacteria indicates that there is much to explore about the ways proteins from other organism interact with their genomes (Hocher et al., 2023). Because it is based on the robust TRIZOL extraction, which has been used on many species across the tree of life (Threlfall and Blaxter, 2021), we expect XDNAX to be applicable to many organisms and tissues. Photo-crosslinking using conventional UV light sources has been successfully reported for the study of protein-RNA interactions in various single-celled organism, as well as embryos of *D.melanogaster*, *C.elegans* and *D.rerio* (Hentze et al., 2018). In a rat xenograft model expanding tumor cells were successfully labeled with 4ST via peritoneal injection (Pridgeon et al., 2011), suggesting that *in vivo* 4ST-labeling of entire animals might be feasible when continuously administered during growth. Preliminary tests with UVEN on mouse tissue slices showed that light penetration of up to one millimeter could be achieved (data not shown), implying that the interrogation of protein-DNA interactions in animal tissue could be feasible.

### Mechanistic insight for the rational design of genotoxic combination therapies

Genotoxic drugs remain among the most applied chemotherapy today, where toxic adverse effects are common and often dose-limiting (Arita et al., 2021). Combination treatments can help to overcome resistance and lower dosage, thereby reducing adverse effects (Al-Lazikani et al., 2012; Lopez and Banerji, 2017). However, a recent screen of clinically relevant drugs in over 2000 combinations found that synergies to decrease the viability of 125 cancer cell lines were rare, with only 5.2 % of all drug combinations and 6.4 % of combinations with a genotoxic drug exhibiting synergy at all (Jaaks et al., 2022). We demonstrate that knockdown of proteins identified on the basis of our DNA interactomes increased the sensitivity of MCF7 cells towards cisplatin, recommending FEN1 or PPP5C inhibition for combination treatments. Indeed, FEN1 has been explored as a potential target for small molecule inhibition before (He et al., 2016, 2017; Tumey et al., 2005; Ward et al., 2017), and treatment of an MCF7 xenograft model with a FEN1 inhibitor in combination with cisplatin synergistically limited tumor size in mice (Dong et al., 2023; Guo et al., 2020). While PPP5C has not been investigated as a drug target by itself, a recent study reported that the small molecule LB-100 inhibits PPP5C as an off-target alongside its designated target PPP2CA (D’Arcy et al., 2019). PPP5C removes inhibitory phosphorylations on ATM and ATR kinase for their efficient activation (Ali et al., 2004; Zhang et al., 2005), implying that its inhibition would forestall recognition of cisplatin-induced DNA damage. LB-100 has been shown to synergize with various DNA-damaging drugs in inhibiting tumor growth in mouse xenograft models (Bai et al., 2014; Lu et al., 2009; Lv et al., 2014) and is currently explored in clinical trials for several cancer indications. Our findings imply that PPP5C inhibition by LB-100 could be a viable way to amplify the genotoxic effect of cisplatin. For future applications of our methodology we envisage that iterative comparisons of DNA interactomes from cells treated with panels of many more chemotherapeutic drugs, and emerging drug combinations, will help the design of increasingly refined genotoxic treatments.

## Acknowledgements

We thank all members of the Kuster laboratory for continuous support and discussion, in particular Karl Kramer, Polina Prokofeva, Johanna Tueshaus and Guillaume Medard. We thank Sara Brajkovic, Nicola Berner, Xintong Sui and Julian Müller for help with the MS measurement. We thank Patroklos Samaras and Tobias Schmidt for help with the initial LED crosslinking tests. We thank Nicholas Kuhn (UCSF) and Christian Frese (Bayer) for advice and discussion. We gratefully acknowledge funding by the German Research Council DFG supporting this work (DFG project number 492625837 and SFB1309-B02 project number 325871075). We also acknowledge funding from the Bavarian research foundation for construction of the UVEN irradiation system (BFS project number AZ-1444-20C).

## Author Contributions

J.T. and B.K. conceived and directed the project and wrote the manuscript. J.T. and S.T. conceived the irradiation device UVEN. S.T. designed and constructed UVEN. J.T. performed *in vitro* benchmarking of UVEN. J.T. performed the XDNAX experiments. J.T. and S.S. performed other proteomic analysis. J.T., S.S. and B.K. measured the proteomic samples. S.S. and J.T. performed knockdown experiments and live cell imaging.

## Supplemental Information

Table S1: Comparison of the direct human DNA interactome to the nuclear and cytosolic proteome of MCF7 cells.

Table S2: Peptides identified with 321 Da modification indicating photo-induced DNA crosslink.

Table S3: Differential quantification of DNA interactomes from MCF7 cells exposed to estrogen for 45 minutes.

Table S4: Differential quantification of DNA interactomes after 24 hour genotoxic stress.

## Methods

### Materials availability

The high-intensity UV irradiation system UVEN is developed as an open science project, which will be described in detail in an upcoming publication. Plans, construction manuals and software for the device can be found at www.uven.org.

### Data Availability

Proteomic data and search results have been deposited in the MassIVE database under the identifier MSV000094079.

### Cell Lines

MCF7 (human [Homo sapiens], female breast adenocacrinoma) and U2OS (human [Homo sapiens], female bone sarcoma) were obtained from ATCC. Cells were maintained in Dulbecco’s Modified Eagle’s Medium (DMEM) supplemented with 10 % dialysed FBS and Pen-Strep (100 U / ml penicillin, 100 mg / ml streptomycin) at 37 °C, 5 % CO_2_. DMEM for SILAC was supplemented with 1 mM L-lysine and 0.5 mM L-arginine of the individual SILAC labels as well as 1.7 mM light L-proline and 2 mM L-glutamine. For full labeling the SILAC label was introduced during six cell passages in DMEM for SILAC. For the dose-dependent analysis of the DNA interactome after estrogen treatment, cells were expanded and treated in DMEM without phenol red as described below. If not further specified labeling with 4-thiothymidine (4ST) occurred over three days at 100 µM concentration in the culture media added from a 100 mM stock in DMSO.

### UV measurements and monitoring of 4SU/4ST photo-conversion by spectrophotometry or LC-MS

Light intensities of the UVEN irradiation system and the reference device VILBER Bio-Link were measured using an LS-128 UV energy meter (Linshang). The sensor was placed behind a conventional polystyrene cell culture dish (Greiner CELLSTAR) at a distance of 3.5 cm from the bulb or LED. Photo-conversion of 4ST and 4-thiouridine (4SU) was monitored in 90 µl of a 1 mM solution irradiated in glass vials at 3.5 cm distance from the bulb or LED. After irradiation the sample was combined with 10 µl of formic acid 1 % and 10 µl analyzed by LC-MS using an Ultimate 1100 HPLC system (Agilent Technologies) connected to an Amazon Speed ETD ion-trap mass spectrometer (Bruker Daltonics). The compounds were resolved across a 30-minute gradient from 10-90 % solvent B (0.1% formic acid in acetonitrile) combined with solvent A (0.1% formic acid in ultrapure water) on a C18 column. The mass spectrometer operated in the positive ion mode with a detection range was of 200 – 1600 m/z. In addition, the absorbance spectrum of the irradiated solutions was recorded on a Nanodrop spectrophotometer (Thermo Scientific).

### Live cell imaging for monitoring proliferation, apoptosis and cell death

Cells were monitored in culture by time-lapse microscopy with an IncuCyte S3 automated microscopy system (Essen Bioscience) in a conventional cell culture incubator. For monitoring proliferation in the presence of 4ST or 4SU, 5000 MCF7 cells or 10000 U2OS cells were seeded onto a 96-well tissue culture plate (testplate 96F, TPP) in 200 µl DMEM and cultured at normal conditions. For investigating the UV sensitization of MCF7 cells after 4ST exposure the 96-well plate was irradiated with UVEN after 4 days of culture. In order to control for potential toxic effects of 4ST or 4SU photo-products in the medium, cells were washed with PBS, and the medium replaced with fresh DMEM without nucleotide before the incubation was continued.

For monitoring the effect of genotoxic drugs 5000 MCF7 cells were seeded and expanded for three days in 100 µl DMEM. Subsequently, 100 µl genotoxic drugs were added at 200 µM concentration (100 µM final concentration) in media containing 63 ng/ml of the apoptosis fluorescence reporter annexin-V CF 594 as well as 100 nM of the cell death fluorescence reporter oxazole yellow homodimer. Because of the incompatibility of platinum compounds with DMSO (Hall et al., 2014), cisplatin and oxaliplatin were directly dissolved to a final concentration of 100 µM in DMEM and etoposide added from a 1:1000 stock in DMSO.

For the monitoring of dose-dependent genotoxic drug effects during siRNA-mediated knockdown a 1:2 dilution series starting from 3.2 mM in 50 µl DMEM was prepared and combined with 50 µl DMEM containing 25 ng annexin-V CF 594 as well as 400 nM oxazole yellow homodimer (final starting concentration 1.6 mM genotoxic drug). This mix was added onto the 100 µl culture medium to yield a final concentration of 800 µM – 12.5 µM of the genotoxic drugs on the cell culture plate.

IncuCyte scans were acquired every two hours using a 4 x objective with phase and additionally green (300 ms) and red (400 ms) channel acquisition if required. For analysis a standard IncuCyte definition was applied using a segmentation adjustment of 1.1 for phase, Top-Hat segmentation for the green channel (100 µM radius, 70 GCU threshold), as well as the red channel (100 µM radius, 1 RCU threshold). For the quantification of proliferation (phase) the confluence (%) and for the quantification of apoptosis (red) or cell death (green) total areas (µM^2^ / image) were exported.

### Knockdown of candidate proteins using siRNA pools

Candidate proteins were knocked down in MCF7 cells with siRNA pools containing 30 distinct siRNAs directed against the target transcript designed and provided by siRNA tools biotech. Knockdown was performed according to the manufacturer’s instructions and efficiency validated by LC-MS. In brief, solution A was prepared by combining 1230 µl of Opti-MEM with 20 µl Lipofectamine RNAiMAX and vortexed vigorously. Solution B was prepared by combining 1050 µl Opti-MEM with 200 µl siRNA dilution (150 nM in ultrapure water). Solution A and B were combined, vortexed and lipid-nucleic acid (LNP) particles allowed to form for 15 minutes. The LNP solution was combined with 1.2 million singularized MCF7 cells in 10 ml DMEM and mixed by pipetting. Subsequently, 10000 cells in 100 µl medium were distributed onto a 96-well plate and allowed to adhere for 15 minutes on the bench before transfer into the IncuCyte. Drug treatment occurred 24 hours after transfection by addition of 100 µl drug concentrate as described above.

### Hypotonic nuclear fractionation and protein cleanup for proteomics using single-pot-solid-phase-enhanced-sample-preparation (SP3)

Reference deep proteomic analysis of the MCF7 total proteome as well as their nuclear and cytosolic subproteomes was performed in quadruplicates. Hypotonic nuclear extraction was applied to generate the cytosolic and nuclear fractions. Per replicate a 15 cm diameter dish of 70 % confluent MCF7 cells was harvested by scraping in to ice-cold PBS and pelleted with 500 g centrifugation for 5 minutes at 4 °C. All subsequent steps were performed on ice. The supernatant was discarded and cells spun down to remove all residual PBS. The pellet was suspended in 2 ml ice-cold nuclear isolation buffer (Tris-Cl 10 mM, KCl 60 mM, MgCl2 1.5 mM, NP40 0.1 %), and pipetted ten times with a 1 ml tip to break the plasma membrane. Cells were incubated for 10 minutes on ice and pipetted again ten times. Nuclei were then pelleted by centrifugation with 2000 g, for 10 minutes at 4 °C. The supernatant containing the cytosolic fraction was transferred to a fresh tube and the nuclei pellet suspended in another 2 ml nuclear isolation buffer for washing. Nuclei were again pelleted with 2000 g, for 10 minutes at 4 °C and taken up in 2 ml nuclear isolation buffer and their integrity checked by microscopy. To create equal conditions between all samples MCF7 cells were taken up in 2 ml nuclear isolation buffer for further processing. Protein content of the samples was determined by BCA and 250 µg protein used for further processing. Volumes were adjusted to 300 µl using nuclear isolation buffer. For protein cleanup a modified version of the SP3 protocol was used (Hughes et al., 2014). To avoid SDS precipitation by potassium proteins were completely denatured with guanidinium thiocyanate (GuTC) and nucleic acids fragmented with trifluoracetic acid (TFA). Therefore, samples were combined with 300 µl SP3 lysis buffer (GuTC 5 M, TFA 200 mM), mixed by vortexing, incubated at 90 °C, 700 rpm shaking for 5 minutes. After addition of 100 µl Tris-Cl pH=7.5 1M samples were homogenized by pipetting and allowed to reach room temperature. For protein aggregation 50 µl of SP3 beads (original slurry) were added, samples vortexed, combined with 1 ml ethanol 100 % and again mixed by inversion. After 15 minutes beads were captured on a magnetic rack, supernatants discarded and beads washed four times with 2 ml EtOH 70%, while attached to the magnet. Protein was digested off the beads overnight in 200 µl trypsin digestion buffer (EPPS 50 mM pH=8, 5 mM DTT, 10 µg trypsin per sample) 37 °C, 700 rpm shaking. Cysteines were alkylated by addition of 6 µl chloroacetamide 550 mM while the incubation was continued for 60 minutes. The beads were collected on a magnet for 5 minutes and the supernatant transferred to a fresh vial. Samples were topped off with 800 µl formic acid 1 % and peptides cleaned up with Sep-Pak C18 cartridges according to the manufacturer’s instructions. Peptides were fractionated at high pH into 96 fractions and pooled to 48 samples as reported before (Bian et al., 2020).

### Treatment of photo-sensitized cells and UV irradiation

For the SILAC-based cataloguing of DNA interactors 1.5 million MCF7 cells were expanded on a 15 cm diameter culture dish for three days in the presence of 100 µM 4ST until approximately 70 % confluent. SILAC light cells were used as unirradiated control and SILAC heavy cells were UV irradiated for protein-DNA crosslinking. Therefore, the medium was decanted, cells washed once with 50 ml ice-cold PBS, covered with 20 ml ice-cold PBS and kept on ice. All subsequent steps were carried out in a cold-room at 4 °C. Photo-crosslinking occurred with the UV irradiation device UVEN for 60 seconds amounting to a combined irradiation energy of 125 J/cm^2^ at 365 nm wavelength. Cells were scraped into the 20 ml ice-cold PBS they were irradiated in pelleted by centrifugation and stored at −80 °C until further use. For the dose-dependent analysis of the DNA interactome in response to estrogen, DMEM without phenol red was used. After three days of expansion in the presence of 4ST MCF7 cells were washed twice with PBS, switched to serum-free DMEM supplemented with 4ST and cultivated for another 48 hours before addition of estrogen (17ß-estradiol) for 45 minutes from a 1:1000 stock in DMSO. Final estrogen concentrations were 10 nM, 3 nM, 1 nM, 300 pM, 100 pM, 30 pM, 10 pM, 3 pM, 1 pM, 300 fM, 100 fM, 30 fM and DMSO control. UV irradiation occurred as described above.

For comparing the DNA interactome of cells treated with genotoxic drugs MCF7 cell were expanded in the presence of 4ST for three days. Consequently, the media was discarded and replaced by fresh media containing the genotoxic drugs or a DMSO mock control alongside 4ST for 24 hours. Because of the incompatibility of platinum compounds with DMSO (Hall et al., 2014), cisplatin and oxaliplatin were directly dissolved to a final concentration of 100 µM in DMEM supplemented with 4ST and etoposide added from a 1:1000 stock in DMSO.

UVEN is developed as a community project aiming to make high-intensity photo-activation in living cells accessible to a wide audience of researchers. Access to detailed plans, construction manuals and software under the creative commons license, is provided upon request and will be presented in an accompanying publication showcasing other UVEN photo-crosslinking applications for proteomics, genomics and transcriptomics.

### Protein-crosslinked DNA extraction (XDNAX) for proteomic quantification of the DNA interactome

Cell pellets from one confluent 15 cm diameter culture dish of MCF7 cells were lysed in 1 ml TRIZOL by pipetting until completely homogenous. Lysates were combined with 200 µl chloroform, mixed by inversion and incubated for 5 minutes at room temperature. Tubes were centrifuged with 12000 g for 10 minutes at 4 °C. The aqueous phase containing RNA was discarded and the interphase containing chromatin transferred to a fresh tube avoiding transfer of organic phase. The interphase was disintegrated in 1 ml recovery buffer (Tris-Cl pH=7.5 50 mM, EDTA 1 mM, SDS 1 %) until completely dissolved. DNA was precipitated by addition of 100 µl NaCl 5 M and 1 ml isopropanol, mixed by inversion, incubated for 15 minutes at −20 °C and spun down for 15 minutes with 20000 g at −11 °C. The supernatant was discarded and the pellet washed with 2 ml ethanol 70 %, centrifuged again for 5 minutes and ethanol removed to completion. For hydration 900 µl ultrapure water (ELGA) was added, the pellet detached from the wall of the tube by vortexing, incubated on a rotating wheel for 30 minutes at 4 °C, and further dissolved by pipetting. For RNA digestion 50 µl Tris-Cl pH=7.5 1 M was added along with 20 µl RNase A (0.5 µg/µl) and 20 µl RNase T1 (0.5 µg/µl) and incubated for 30 minutes at 37 °C, 700 rpm shaking. After addition of 5 µl SDS 20 % clumps were dissolved by pipetting and incubation continued for another 30 minutes. For DNA fragmentation one sample was distributed onto one column (8 wells) of a Covaris plate in 120 µl portions and sonicated using the parameters 50 cycles, Scan Speed: 1.0, PIP: 300, CPB: 50, AIP: 75, Dithering: Y=1 Speed=10. The sonicated sample was collected in a fresh tube and centrifuged with 5000 g for 5 minutes at room temperature. The supernatant was transferred to a fresh falcon tube, and the remaining pellet dissolved in sonication buffer (Tris-Cl 50 mM, SDS 0.1 %). Sonication and centrifuging was repeated until no pellet remained. The SDS concentration in the sonicated supernatants was adjusted to 2 % and the sample incubated at 95 °C for 5 minutes, 700 rpm shaking. The sample volume was doubled by addition of guanidinium thiocyanate (GuTC) 5 M, the sample mixed by vortexing and again incubated at 95 °C for 5 minutes, 700 rpm shaking. The volume was doubled again by addition of ethanol 100 %, the sample thoroughly mixed by vortexing and allowed to reach room temperature. The sample was successively applied to a silica spin column (Zymo-Spin IIICG Column) in 800 µl increments using a table top mini centrifuge with approximately 2000 g. We note here that overloading will impair removal of non-crosslinked protein and recommend to stop loading as soon as the column flow decreases notably. Columns were washed twice with 800 µl wash solution (GuTC 2.5 M, ethanol 50 %), and three times with 800 µl ethanol 70 % each time centrifuging 2 minutes with 10000 g. For elution of protein-DNA complexes column was transferred to a fresh tube and the drain hole blocked with parafilm. After addition of 200 µl nuclease elution mix (NEB nuclease P1 buffer 1 x, MgCl_2_ 5 mM, 0.5 µl NEB nuclease P1, 0.5 µg benzonase) samples were incubated over night at 37 °C, 700 rpm shaking. After addition of 20 µl SDS 20 % the incubation was continued for 30 minutes. The sample was recovered into a fresh tube by centrifugation with 16000 g for 5 minutes. Remaining DNA was eluted two more time with 200 µl elution buffer (Tris-Cl 50 mM, SDS 2 %) and the eluates combined. For protein cleanup 10 µl SP3 beads (original slurry) were added and vortexed, resulting in an approximately 600-640 µl of total sample volume. For protein aggregation 1 ml ethanol 100 % was added, the sample mixed by inversion and incubated for 15 minutes without shaking. The beads were collected on a magnet for 5 minutes and the supernatant discarded. Remaining on the magnet, the beads were washed four times with 2 ml ethanol 70 % and subsequently spun down to remove all residual ethanol. Protein was digested off the beads overnight in 200 µl trypsin digestion buffer (EPPS 50 mM pH=8, 5 mM DTT, 0.5 µg trypsin per sample) 37 °C, 700 rpm shaking. Cysteines were alkylated by addition of 6 µl chloroacetamide 550 mM while the incubation was continued for 60 minutes. The beads were collected on a magnet for 5 minutes and the supernatant transferred to a fresh vial. Resulting peptides are heavily contaminated with nucleotides and a polymer of unknown origin (probably PEG from spin columns), which can be removed by SCX StageTip but not C18. We therefore first cleaned up samples by SCX StageTip and then used an additional C18 cleanup for high-pH fractionation. For SCX StageTip 10 µl formic acid 10 % was added along with 100 µl acetonitrile (approximately 30 % final). SCX StageTips were preconditioned with 200 µl acetonitrile and 200 µl SCX wash solution (formic acid 0.1 %, ACN 30). Samples were loaded onto SCX StageTips and washed five times with 200 µl SCX wash solution. Peptides were eluted twice with 50 µl high-pH elution buffer (ammonium formate pH=10 250 mM, 50 % ACN) and dried by SpeedVac. For high-pH fractionation C18 StageTips were preconditioned with 200 µl ACN and 200 µl formic acid 0.1 %. Dried samples were applied in 200 µl formic acid 0.1 % and washed twice with 200 µl formic acid 0.1 %. Fractionation occurred with high-pH buffer (ammonium formate pH=10 50 mM) supplemented with 0, 5, 10, 15, 20, and 50 % acetonitrile. Following fractions were combined and dried down before LC-MS analysis, F1 5 + 50 %, F2 10 %, F3 15 %, F4 0+20 % acetonitrile.

### HPLC-MS for the detection and quantification of proteomic samples

Peptide analysis occurred on Orbitrap Eclipse Tribrid mass spectrometers (Thermo Scientific), connected to a Dionex UltiMate 3000 RSLCnano system (Thermo Scientific) for nanoflow detection of XDNAX samples, or to a Vanquish Neo UHPLC system (Thermo Scientific) for microflow detection of 48-fraction deep reference proteomes.

For nanoflow analysis, the sample was injected onto a trap column (75 μm x 2 cm) packed with 5 μm C18 resin (Dr. Maisch Reprosil PUR AQ) in solvent C (formic acid 0.1 %). Peptides were washed with solvent C at 5 µl/min for 10 minutes and subsequently transferred on to an analytical column (75 μm x 48 cm, heated to 55 °C) packed with 3 μm C18 resin (Dr. Maisch Reprosil PUR AQ) using a gradient of solvent A (formic acid 0.1 % in ultrapure water, DMSO 5%) and solvent B (formic acid 0.1 % in acetonitrile, DMSO 5%). Elution occurred across a 60-minute gradient at a flow rate of 300 nl/min starting from 4 % B followed by a linear increase to 32 % B. Nanosource voltage was 2000 V, ion transfer tube temperature 275 °C. Detection occurred with data-dependent acquisition using an OT-OT method and a cycle time of 2 seconds. MS1 resolution was 60000, scan range 360-1300, RF lens 40 %, AGC target 100 % and maximum injection time 50 ms. MS2 isolation occurred with a quadrupole window of 1.2 m/z and fragmentation with 30 % HCD energy. MS2 resolution was 30000, first mass 100 m/z, AGC 200 % and maximum injection time 54 ms.

For microflow analysis, the sample was directly injected onto an Acclaim PepMap 100 analytical column (2 µm particle size, 1 mm x 150 mm, heated to 55 °C) for 1.4 s at 100 µl/s flow of solvent A (formic acid 0.1 % in ultrapure water, DMSO 3 %) combined with 5.2 % of solvent B (formic acid 0.1 % in acetonitrile, DMSO 5%). Elution occurred at a flow rate of 50 µl /min starting from 5.2 % B, followed by a linear increase to 23 % B until 12.4 minutes, followed by a linear increase to 28% B until 13.6 minutes. Source voltage was 3500 V, ion transfer tube temperature 325 °C, vaporizer temperature 125 °C. Detection occurred with data-dependent acquisition using an OT-OT method and a cycle time of 0.9 s. MS1 resolution was 120000, scan range 360-1300, RF lens 40 %, AGC target 100 % and maximum injection time 50 ms. MS2 isolation occurred with a quadrupole window of 1.3 m/z and fragmentation with 28 % HCD energy. MS2 resolution was 15000, first mass 100 m/z, AGC 200 % and maximum injection time 22 ms.

### MS database search

MS raw files were searched using MaxQuant. In addition, MSfragger was used to identify 4ST-modified peptides.

All MaxQuant settings were used at their default value, except for specifying SILAC configurations and indicating the appropriate number of fractions per sample. Label-free quantification with iBAQ was activated and used at default parameters. In case of the differential quantification of DNA interactomes from MCF7 cells treated with estrogen or genotoxic drugs the ‘match between runs’ option was activated and used at default parameters. In case of the differential quantification of DNA interactomes from MCF7 cells treated with estrogen, as well as the SILAC-based quantification of four-hour etoposide treatment ‘phospho STY’ was added as additional variable modification. In these two cases the MaxQuant search was performed with default FDR settings and standard proteinGroups.txt and Phospho(STY)Sites.txt tables used for further analysis. In case of the MCF7 DNA interactome, the deep reference proteomes and the quantification of DNA interactomes in response to genotoxic drugs the MaxQuant search was performed 100 % FDR followed by PROSIT rescoring (Gessulat et al., 2019). Therefore, ‘protein FDR’ and ‘peptide FDR’ were set to 1 and the search results rescored by an in-house pipeline of PROSIT followed by picked-FDR thresholding (The et al., 2022).

MSfragger was used as part of FragPipe. Raw files for the SILAC-controlled MCF7 DNA interactome were first surveyed with an open search using parameters preset in the software. Precursor mass tolerance was set to −150 – 1000 Da, add_C_cystein was set to 57.021464, add_R_arginine was set to 10.0083, add_K_lysine was set to 8.0142, and the export format set to TSV_PEPXML_PIN. In the validation section Crystal-C was activated, and PSM validation set to the recommended open search parameters. In the FDR filter and report section ‘Generate peptide-level summary’ was selected. All DNA interactome data was also searched for the 321 Da phospho-4ST-H_2_O modification in an offset search with parameters at their default values. Mass offsets were set to ‘0 321.0316’, and the modification add_C_cystein to 57.021464, as well as add_R_arginine to 10.0083, add_K_lysine to 8.0142 for the SILAC-controlled experiment. In the validation section Crystal-C was de-activated, and PSM validation set to the recommended offset search parameters.

### Processing and analysis of proteomic data

All proteomic data was processed in R(4.2.2) using RStudio. For the analysis of proteomic data rescored with PROSIT the PickedFDR proteinGroups_fdr0.01.txt file was used. MaxQuant contaminants were filtered by removing all ‘CON’ entries. In case of the standard MaxQuant searches including the phospho-STY modification the proteinGroups.txt table was used and filtered to remove ‘Potential contaminants’, ‘Reverse’ matches to the decoy database, as well as proteins ‘Only identified by site’. In order to determine the MCF7 DNA interactome each SILAC-controlled replicate of was processed independently. The aim of our enrichment strategy was to eliminate as much non-crosslinked protein (SILAC light) and recover as much DNA-crosslinked protein as possible (SILAC heavy). As a proxy for protein quantities in each SILAC channel iBAQ values were used (Schwanhäusser et al., 2011). Proteins with only iBAQ intensity in the irradiated SILAC channel were immediately called as DNA interacting in this replicate and proteins with only iBAQ intensity in the unirradiated SILAC channel as non-crosslinked protein. For all other proteins iBAQ rations of log2(irradiated (SILAC heavy) / unirradiated (SILAC light)) were calculated and the apex of the resulting distribution determined to account for mixing errors between the SILAC channels (Figure S3A). The left side of this distribution was then mirrored to create a symmetrical lognormal distribution estimating iBAQ ratios among non-crosslinked proteins. Three standard deviations from the apex of this mirrored distribution were used a replicate-specific cutoff to call DNA interacting proteins. Proteins called DNA interacting in at least three out of five replicates were included in the MCF7 DNA interactome (Table S1).

Deep reference proteomes were analysed to determine the MCF7 nuclear and cytosolic proteome using iBAQ quantification, which were normalized by median centering between the four replicates of one experiment. As illustrated in Figure S3D the imperfect hypotonic nuclear fractionation and our exhaustive peptide fractionation strategy led to the detection of most MCF7 proteins contained in the full proteome also in the cytosolic fraction, albeit, at very different abundance (Table S1). Because we were focused on the proteomic surrounding of DNA we defined the nuclear proteome using a quantitative cutoff oriented on transcription factors. Figure S3C shows that the iBAQ ratio nucleus / cytosol for transcription factors was in 89 % of cases higher 0.1. The remaining 11 % contained mostly transcription factors know to be naturally localized to the cytosol, such as STAT3, IRF3, AR, NFKB1, RELA etc., so that we used a cutoff of 0.1 to call proteins present in the nucleus. Vice versa, proteins with an iBAQ ratio smaller 0.1 were defined as cytosolic.

For the differential analysis of the single-dose estrogen treatment as well as the genotoxic drug treatments we used iBAQ quantification. Under the prerequisite that iBAQ values represent counts of protein molecules we applied negative binomial model-based differential analysis with DESeq2 (Love et al., 2014). Foldchanges were corrected using the apeglm package (Benhalevy et al., 2018). Because of the extreme changes in the DNA interactome of cells treated with genotoxic drugs, where many interesting proteins such PSMC2, SELENOH, CREB1, TP53, CHEK2 etc. (Table S4) were undetected in one of the conditions, we applied imputation of missing values in order to include these proteins in our differential analysis. Therefore, we used an adaptation of the imputation function described for PERSEUS (Tyanova et al., 2016), which draws random numbers from a down-shifted normal distribution with shrunken standard deviation using the parameters width 0.3 and downshift 1.8.

For dose-response analysis of DNA interactomes in response to estrogen iBAQ quantification was used (Table S3). Therefore protein abundances from all DNA interactomes in the analysis were first normalized by median centering. To avoid overfitting quadruplicates of the highest estrogen concentration and the DMSO control were collapsed into one replicate each using their median. For fitting a log-logistic model the R package ‘drc’ was used.

### Analysis of live cell imaging data

Growth inhibition by the photo-activatable nucleotides 4ST and 4SU towards MCF7 or U2OS cells was analyzed by summing up the confluence data recorded over five days for each individual replicate of each concentration. The dose-dependent induction of apoptosis or cell death by cisplatin (Figure 6C&D) was analysed in the same way by summing up fluorescence measurements for annexin-V or oxodazol yellow (see above) over the 3 days of the knockdown experiment. The R package ‘drc’ was used to fit a log-logistic dose-reponse model and derive effective concentrations.

### Functional annotation of proteins and data visualization

For all cross-references with protein databases UniProt identifiers were used, except for OMIM where gene names were used. Proteins were annotated with their gene ontology (GO) terms via ENSEMBL BioMart accessed via the R package ‘biomaRt’. We note here that for clarity GO was used as only source to annotate proteins as ‘DNA binding’ or ‘RNA binding’. In some cases this varies from their annotation in UniProt, which integrates annotation from different sources. GO enrichment analysis was performed using the GOrilla web interface. Protein structures were visualized in UCSF Chimera. Data was plotted using the R package ‘ggplot2’. Schemes in Figures 2A and 3J were created in part with Biorender.

---

**Figure S1:**
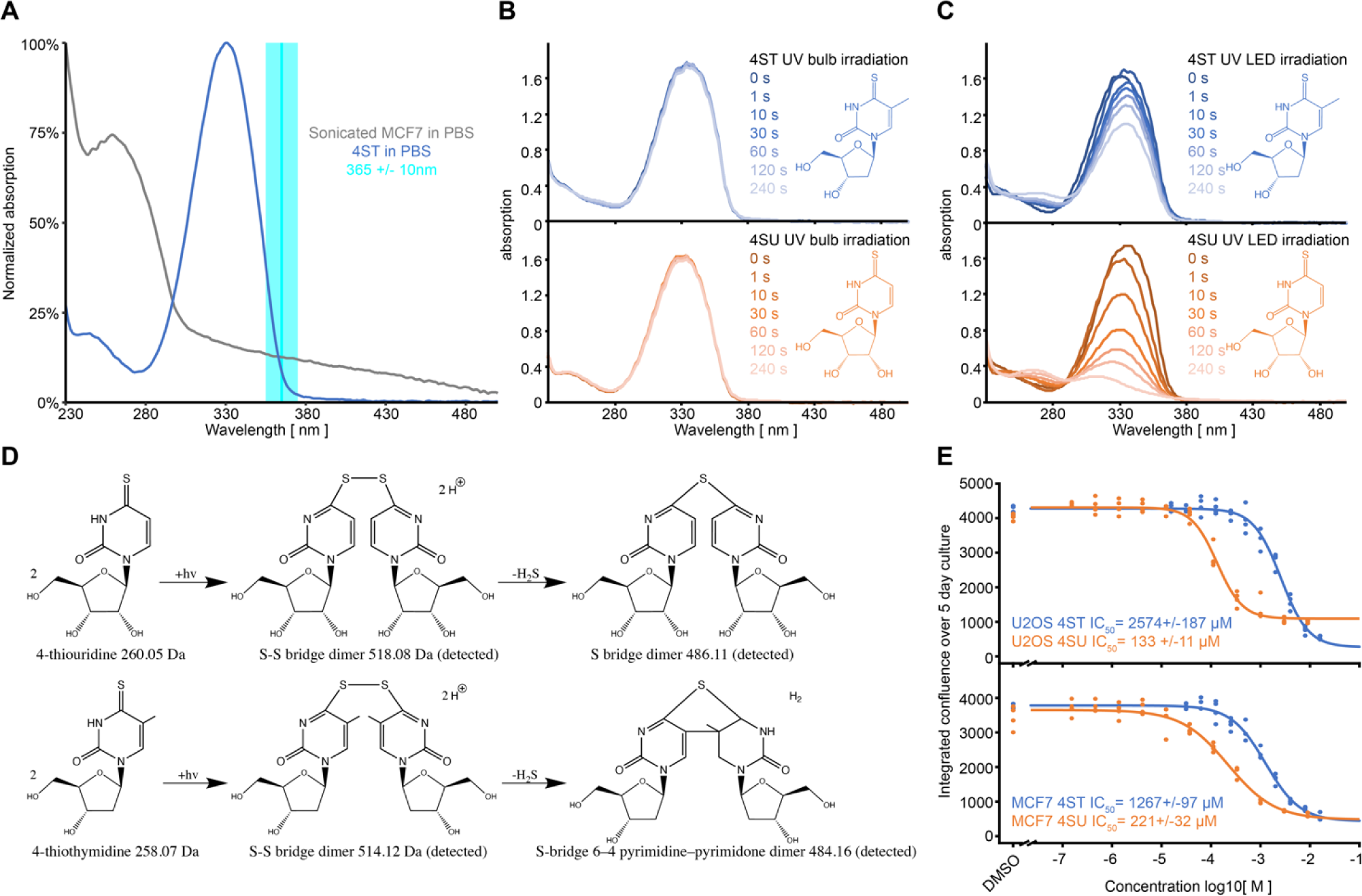
Photo-activation of 4-thiothymidine and metabolic labelling of living cells. A) Lineplot comparing the light absorption of 4ST in PBS to MCF7 cells sonicated to homogeneity in PBS. B) Absorption spectra of 4ST (top) and 4SU (bottom) irradiated for indicated time with a conventional UV bulb irradiation device. See also Figure 1D. C) Same as in B but irradiation with UVEN. See also Figure 1D. D) Reaction scheme for the formation of photo-dimers from 4ST (top) or 4SU (bottom). Reaction products detected by LC-MS are indicated. E) Dose-response analysis for the growth inhibition of U2OS (top) or MCF7 cells (bottom) by 4ST (blue) or 4SU (brown) added to the culture medium. Cell confluence was monitored by live cell imaging and integrated over 5 days of culture. Points indicate individual replicates and the line a log-logistic model.

**Figure S2:**
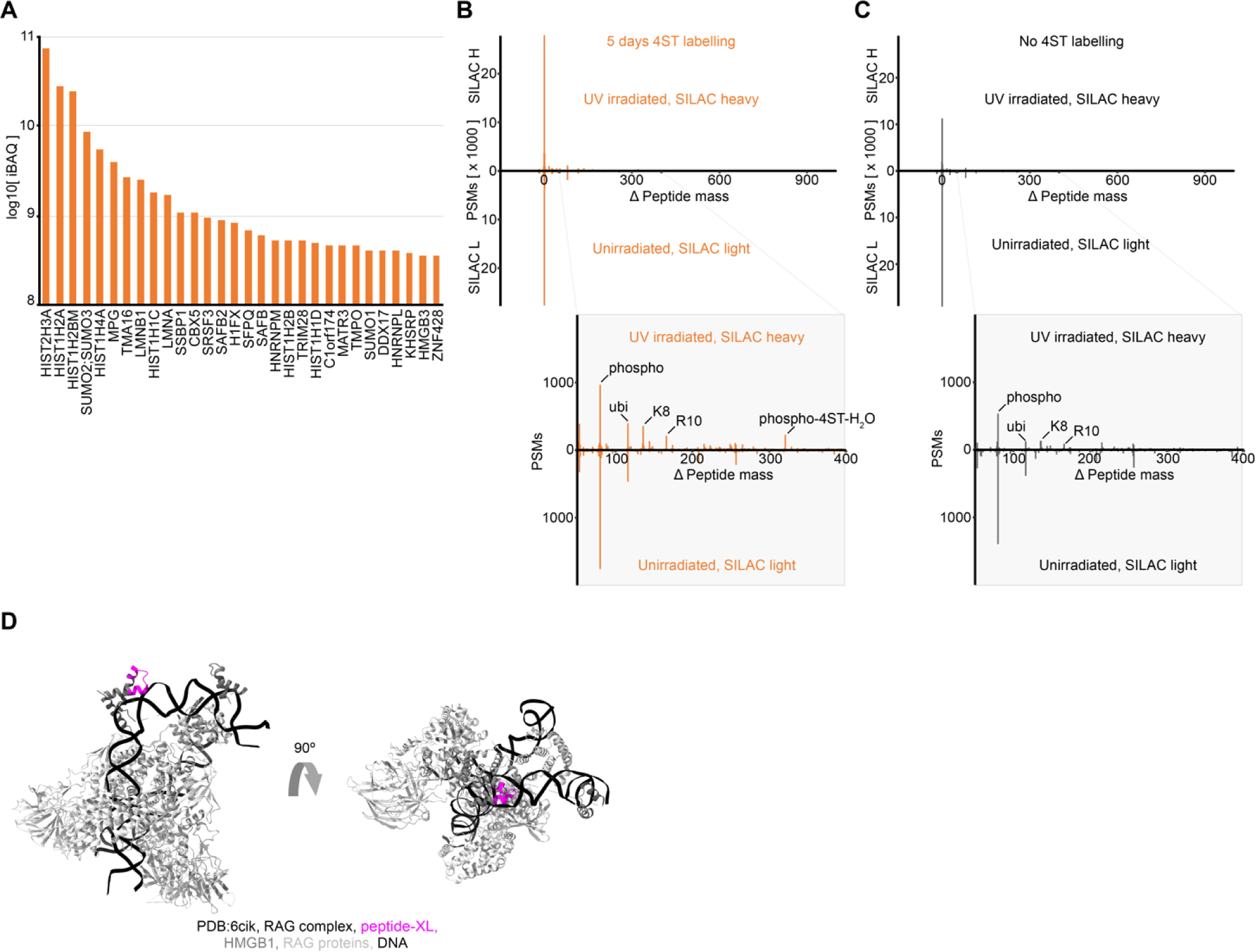
Most abundant proteins and nucleotide-peptide hybrids extracted by XDNAX. A) Bar plot illustrating the most abundant proteins identified in protein-crosslinked DNA enriched by XDNAX. Displayed are the top 30 proteins from the irradiated SILAC channel of Figure 2C ranked by iBAQ. B) Histogram of peptide-spectrum matches (PSMs) identified in an open search by the mass-tolerant search engine MSfragger. The magnified area highlights the occurrence of mass adducts corresponding to protein phosphorylation (phospho), uniquitination (ubi), heavy lysine and arginine (K8, R10, SILAC artefacts), as well as a new modification corresponding to 4ST with a water loss (4ST-H_2_O). C) Same as B but for cells without 4ST labelling. See Figure 2D. D) Crystal structure of the RAG1 pre-reaction complex containing HMGB1 (PDB 6CIK). The nucleotide-crosslinked peptide from HMGB1 identified in our DNA interactome (magenta) localizes to the direct vicinity to the DNA (black).

**Figure S3:**
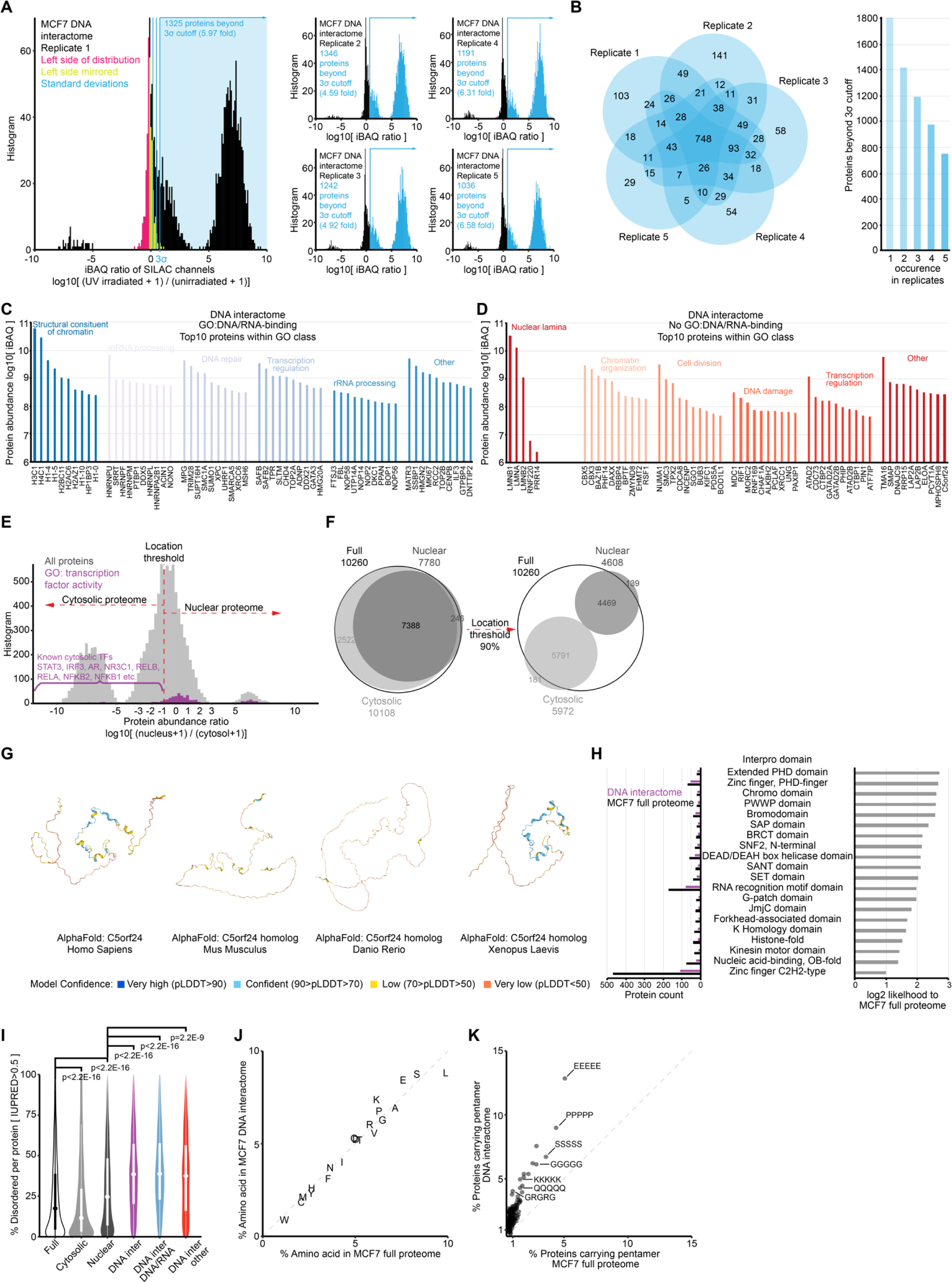
Defining the direct DNA interactome as well as the nuclear, cytosolic and full proteome of MCF7 cells using quantitative proteomics. A) Histograms showing the filtering process for the generation of the direct DNA interactome from five replicates of photo-crosslinked MCF7 cells subjected to XDNAX. For each replicate a background distribution was estimated by mirroring negative log-foldchanges under the assumption that non-crosslinked proteins exhibit the same intensity variance in both SILAC channels. We used three standard deviations (3σ) of this distribution as fold-change threshold for calling candidate DNA-interacting proteins in each replicate. B) Venn diagram showing the occurrence of candidate DNA-interactors for each replicate (left), and barplot comparing occurrence of proteins across replicates (right). For the final DNA interactome shown in Figure 3A and used throughout the rest of our analysis, only candidate DNA-interacting proteins occurring in three or more replicates were considered. C) & D) Bar plot showing the top ten proteins contributing the largest iBAQ to each GO term in Figure 3A. Shown are means of the iBAQ from the SILAC channel of photo-crosslinked cells (SILAC heavy). E) Histogram of protein abundance ratios between the MCF7 nuclear and cytosolic proteome. An arbitrary abundance cut-off was chosen to include as many transcription factors in the nuclear proteome as possible, while excluding transcription factors know to be constitutively located in the cytosol (STAT3, IRF3, AR, NR3C1, RELB, RELA, NFKB2, NFKB1 etc.). For details see Methods. F) Venn diagrams showing proteins contributing to the nuclear and cytosolic proteome of MCF7 cells before (left) or after (right) applying the abundance threshold established in E. Because the focus of this study was the nucleus we considered all proteins part of the nuclear proteome that contributed more than 10 % of their iBAQ to the nuclear fraction. For simplicity the cytosolic proteome was defined by exclusion of nuclear proteins. For details see Methods. G) Protein structure predictions by AlphaFold for the human C5orf24 and its homologues in mice, zebra fish and frogs. H) Barplots comparing the occurrence (left) and relative likelihood (right) of interpro domains in the DNA interactome relative to the MCF7 full proteome. I) Violin plots comparing the percentages of amino acid positions predicted as disordered according to their IUPRED score (IUPRED>0.5) (Erdos et al., 2021). Testing occurred with a two-sided Kolmogorov-Smirnov test. Proteins in the MCF7 full proteome (full) are compared to the cytosolic and nuclear proteome (see Figure S3F), the DNA interactome (DNA inter) and its parts annotated as nucleic-acid binding (DNA/RNA) or lacking this annotation (other). J) Scatter plot comparing the amino acid frequency in proteins of the MCF full proteome to the DNA interactome. K) Scatter plot comparing the occurrence of disordered amino acid pentamers in proteins of the MCF7 full proteome to the DNA interactome. All possible pentameric permutations of the amino acids G, S, D, Q, P, E, K and R were counted if they occurred at least once in a protein in the two groups.

**Figure S4:**
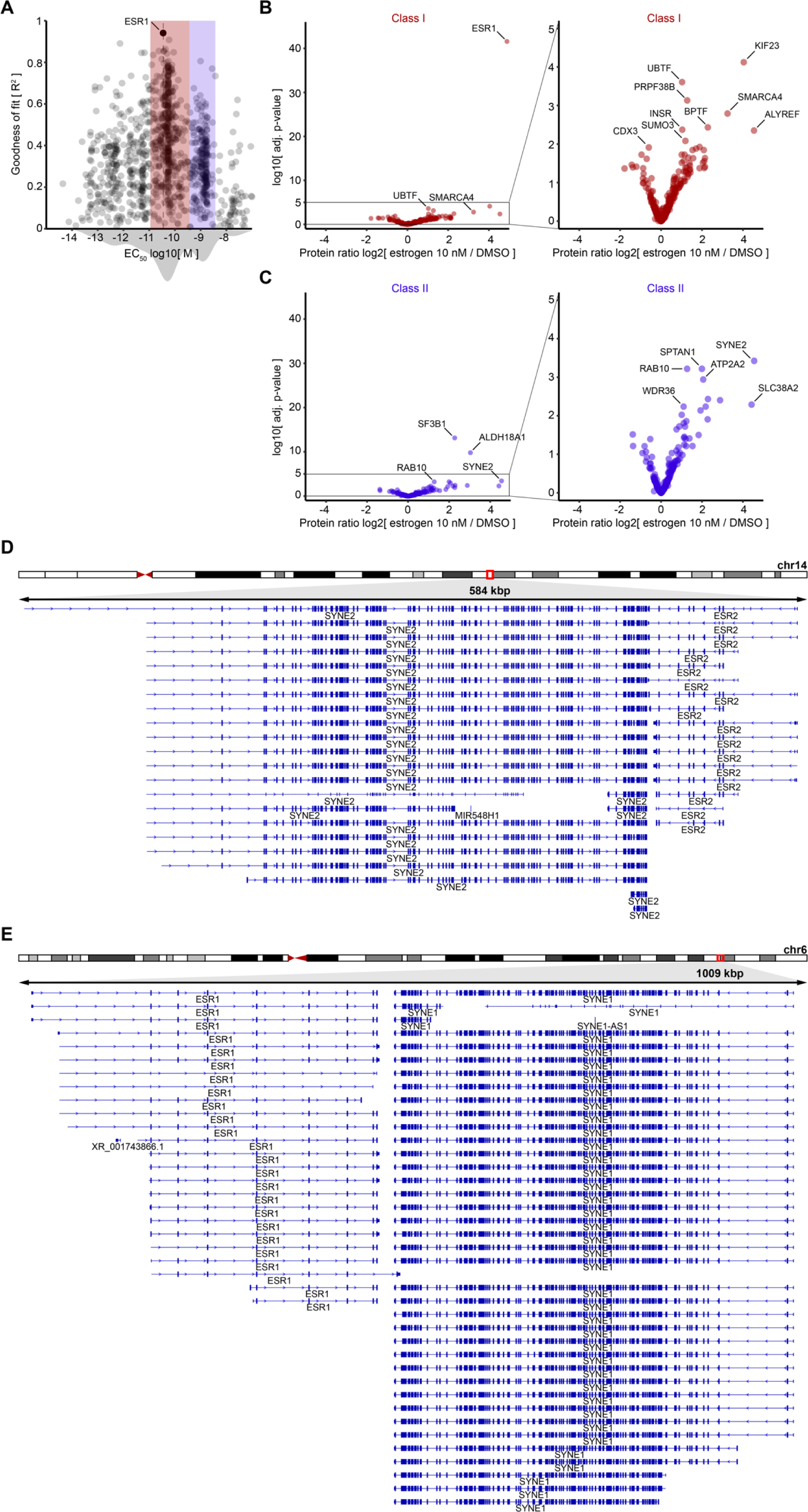
Primary and secondary changes in protein-DNA interactions during estrogen exposure. A) Scatter plot comparing the half-effective concentration (EC_50_) and the goodness of fit (R^2^) for 1814 proteins in the DNA interactome of MCF7 cells exposed to 13 concentrations of estrogen for 45 minutes. B) Volcano plot illustrating changes in the DNA interactome during 45 minutes of high-dose estrogen exposure only showing class I proteins identified in A. C) Volcano plot illustrating changes in the DNA interactome during 45 minutes of high-dose estrogen exposure only showing class II proteins identified in A. D) The human ESR2 locus on chromosome 14 with the neighbouring SYNE2 gene. Refseq annotations of transcript isoform are given in blue. E) The human ESR1 locus on chromosome 6 with the neighbouring SYNE1 gene. Refseq annotations of transcript isoform are given in blue.

**Figure S5:**
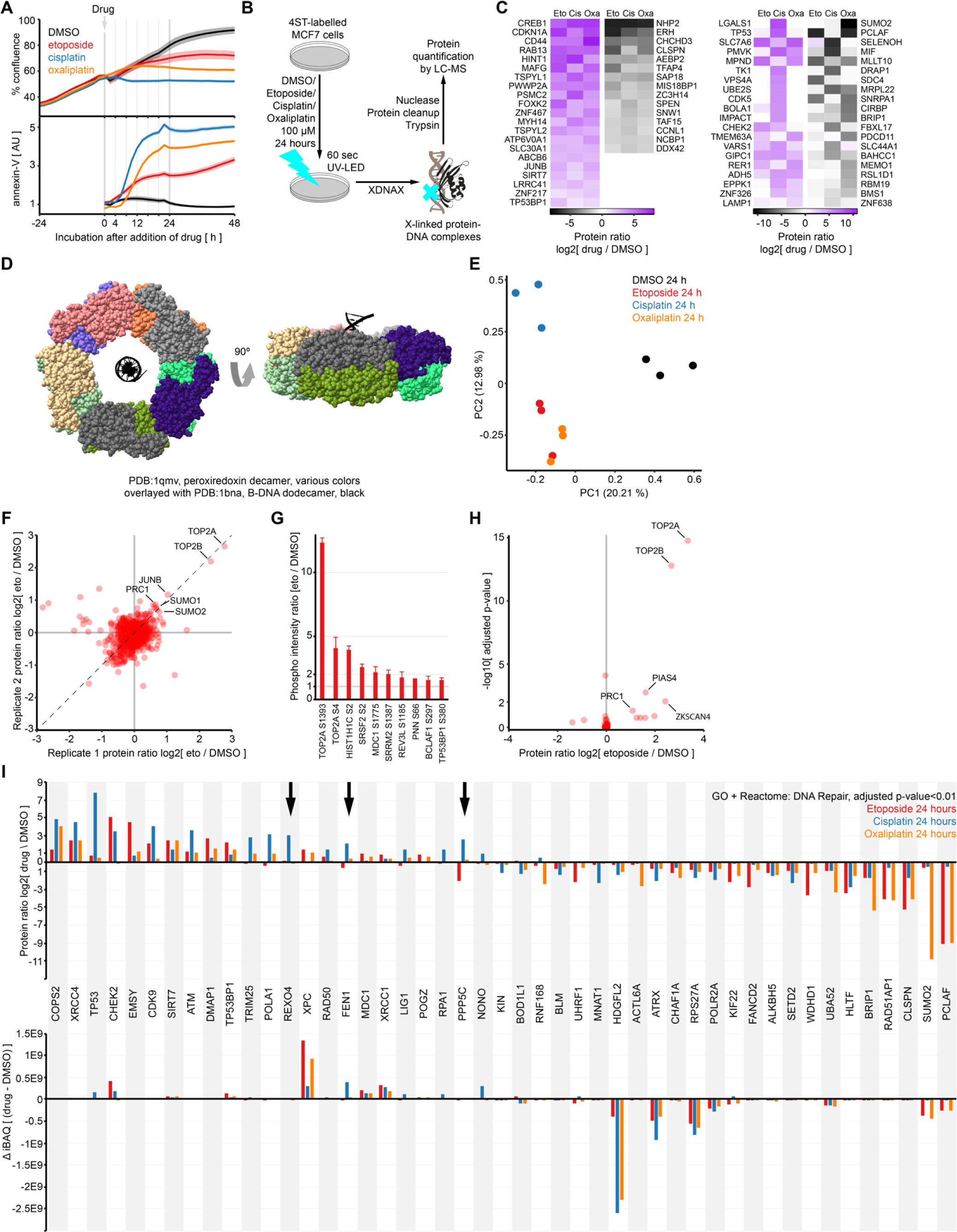
Changes in the direct DNA interactomes from breast cancer cells exposed to different genotoxic drugs. A) Lineplot showing proliferation (top, confluence) and induction of autophagy (bottom, Annexin-V flouresence) monitored by live cell imaging. After 24 hours of normal expansion genotoxic drugs were added at 100 µM concentration. Shown are the mean of ten replicates as bold line and their standard error as shading. B) Experimental outline for the comparison of proteins interacting with DNA after 24 hours of treatment with a genotoxic drug or mock control (DMSO). C) Left: Heatmap displaying relative abundance ratios of all proteins with significantly changed DNA interaction across all three genotoxic drugs compared to the mock control (adj. p<0.01 in all three treatments). Right: Heatmap displaying relative abundance ratios of proteins with significantly changed DNA interaction in maximal two treatments (adj. p<0.01 not in all three treatments). The top 20 proteins with the most extreme increase (left) and decrease (right) are displayed. D) Protein structure overlay of the peroxiredoxin II decamer (PDB 1qmv) and a DNA twelve-mer (PDB 1bna) visualizing that by proportion the DNA helix could be encompassed by a PRDX2 toroid. E) PCA analysis of the triplicate DNA interactomes from MCF7 cells using all protein abundances (iBAQ) as input. Missing values were imputed to include extreme changes (e.g. TP53, CREB1, PCLAF etc., see Table 4). F) Scatter plot comparing normalized SILAC ratios between replicates of DNA interactomes from MCF7 cells treated with 100 µM etoposide for 4 hours or a DMSO control. SILAC labels were swapped between the replicates. G) Barplot showing intensity ratios for ten phospho peptides in B with the most extreme foldchanges. Error bars indicate composite standard deviations from three four replicates. H) Volcano plot illustrating changes in the DNA interactome after four hours of etoposide treatment. Four replicates of differentially treated SILAC samples were used. I) Barplot comparing relative protein abundance changes (top) to absolute abundance changes (bottom) in DNA interacomtes from gentoxic drug-treated cells compared to mock-treated cells. Displayed are all proteins that tested significant (adj. p<0.01) in at least one of the conditions, implying that some of the indicated values do not represent significant changes (see Table S4). Arrows indicate potential cisplatin sensitivity factors chosen for follow-up experiments (see Figure 6).

**Figure S6:**
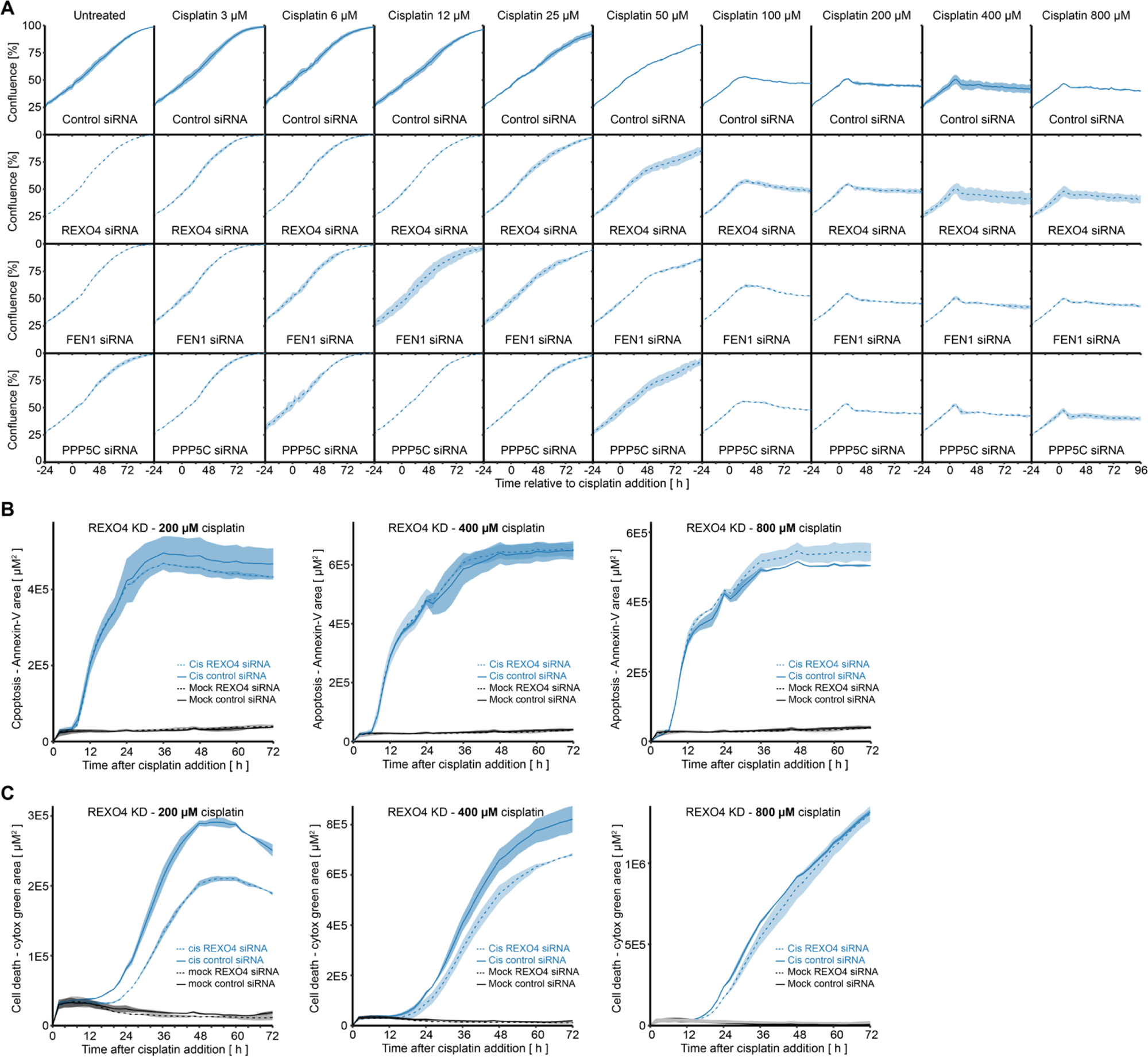
Live cell imaging time series confirming DNA interactors as cisplatin sensitivity factors. A) Lineplots following the proliferation of MCF7 cells with knockdowns for REXO4, FEN1 or PPP5C using automated microscopy. One uniform pool of cells was transfected with siRNAs 24 hours before addition of the cisplatin concentration indicated on top. B) Lineplots following the induction of apoptosis in MCF7 cells with knockdowns for REXO4 using fluorescence-labelled Annexin-V. One uniform pool of cells was transfected with siRNAs 24 hours before addition of 100, 400 or 800 µM cisplatin. Bold lines indicate cells transfected with control siRNA, dotted line for REXO4 siRNA, shading indicates one standard deviation. C) Same as in B but for the quantification of dead cells using the fluorescence dye cytotox green.

